# Epithelial GPR35 protects from *Citrobacter rodentium* infection by preserving goblet cells and mucosal barrier integrity

**DOI:** 10.1101/2021.03.27.437264

**Authors:** Hassan Melhem, Berna Kaya, Tanay Kaymak, Philipp Wuggenig, Emilio Flint, Julien Roux, Claudia Cavelti-Weder, Maria L. Balmer, Jean-Claude Walser, Rodrigo A. Morales, Christian U. Riedel, Prisca Liberali, Eduardo J. Villablanca, Jan Hendrik Niess

**Affiliations:** Department of Biomedicine, Gastroenterology, University of Basel, 4031 Basel, Switzerland; Swiss Institute of Bioinformatics, 4031 Basel, Switzerland; Department of Biomedicine, Endocrinology, Diabetes, and Metabolism, University Hospital of Basel, 4031 Basel, Switzerland; Department of Diabetes, Endocrinology, Nutritional Medicine and Metabolism, Bern University Hospital, University of Bern, 3010 Bern, Switzerland and Diabetes Center Berne, 3010 Bern, Switzerland; Genetic Diversity Centre, Department of Environmental Systems Sciences, ETH Zurich, 8092 Zurich, Switzerland; Division of immunology and Allergy, Department of Medicine, Solna, Karolinska Institute and University Hospital, 17176 Stockholm, Sweden; Center of Molecular Medicine, 17176 Stockholm, Sweden; Institute of Microbiology and Biotechnology, University of Ulm, Ulm, Germany; Friedrich Miescher Institute for Biomedical Research, Basel, Switzerland; University of Basel, Basel, Switzerland; University Center for Gastrointestinal and Liver Diseases, St. Clara Hospital and University Hospital of Basel, 4031 Basel, Switzerland

## Abstract

Goblet cells secrete mucin to create a protective mucus layer against invasive bacterial infection and are therefore essential for maintaining intestinal health. However, the molecular pathways that regulate goblet cell function remain largely unknown. Although GPR35 is highly expressed in colonic epithelial cells, its importance in promoting the epithelial barrier is unclear. In this study, we show that epithelial Gpr35 plays a critical role in goblet cell function. In mice, cell type-specific deletion of *Gpr35* in epithelial cells but not in macrophages results in goblet cell depletion and dysbiosis, rendering these animals more susceptible to *Citrobacter rodentium* infection. Mechanistically, scRNA-seq analysis indicates that signaling of epithelial Gpr35 is essential to maintain normal pyroptosis levels in goblet cells. Our work shows that the epithelial presence of Gpr35 is a critical element for the function of goblet cell-mediated symbiosis between host and microbiota.

## Introduction

Goblet cells are the most abundant secretory epithelial cells in the colon. Their principal functions involve the production and secretion of mucins, thereby providing a thick mucus layer covering the apical surface of the intestinal epithelium. This mucus layer acts as the first line of defense by fending off luminal bacteria, thus reducing bacterial exposure of epithelial and immune cells. Gel-forming O-linked glycosylated Muc2 polymers are the main component of the intestinal mucus and play a crucial role in maintaining a regular microbial community in the gut (Wu et al., 2018). Mucus layer impairment leads to infection and inflammation, as described for inflammatory bowel disease (IBD) (Cornick et al., 2015, Wells et al., 2017). Indeed, ulcerative colitis (UC) has been associated with a reduced number of goblet cells, defective production and secretion of mucins, and increased bacterial penetration (van der Post et al., 2019). *Muc2*-deficient mice display excessive bacterial contact with their colonic epithelium and spontaneously develop chronic colitis (Johansson et al., 2011a, Johansson et al., 2011b, Johansson et al., 2008, Zarepour et al., 2013). The lack of *Muc2* also impairs clearance of the attaching and effacing (A/E) pathogen *Citrobacter rodentium* (*C. rodentium*) (Bergstrom et al., 2010). Intriguingly, the precise mechanisms that alter the mucus layer leading to defective barrier integrity remain largely unknown.

Supporting the hypothesis that the microbiota and their metabolites strongly contribute to the modulation of the intestinal mucus layer which appears thinner in germ-free mice than in conventionally housed mice (Johansson et al., 2015). The microbiota-mediated establishment of intestinal barrier integrity is dependent on signaling through G protein-coupled receptors (GPCRs) (Melhem et al., 2019, Tan et al., 2017). Genome-wide association studies on GPR35 single nucleotide polymorphisms indicated that the rs3749171 variant of GPR35, responsible for T108M substitution, might be related to the pathogenesis of UC (Ellinghaus et al., 2013, Imielinski et al., 2009). GPR35 remains an orphan GPCR, although we recently demonstrated that lysophosphatidic acid (LPA) is a potential endogenous ligand for GPR35 using cell-based assays (Kaya et al., 2020). Besides LPA, several other candidates, including the tryptophan metabolite kynurenic acid (Wang et al., 2006) and the chemokine CXCL17 (Maravillas-Montero et al., 2015) can act as potential endogenous ligands for GPR35. In humans and rodents, GPR35 is expressed by macrophages. However, its expression is particularly prominent in epithelial cells (ECs) (Lattin et al., 2008) suggesting that GPR35 may play a key role in maintaining the integrity of the epithelial compartment. In line with this, GPR35 signaling has been shown to be essential for epithelial cell turnover, renewal, and wound healing in mouse models (Schneditz et al., 2019, Tsukahara et al., 2017). In agreement with these findings, deletion of GPR35 aggravated dextran sulfate sodium (DSS)-induced experimental colitis in mice (Farooq et al., 2018). These observations are highly suggestive of a crucial role of GPR35 in the regulation of epithelial barrier integrity. Nevertheless, the mechanisms by which the IBD risk gene *GPR35* modulates the gut epithelial barrier are still not known.

Colonic ECs are constantly renewed to maintain an intact mucosal barrier (van der Flier and Clevers, 2009). High cellular turnover is critically regulated by different modes of cell death including pyroptosis (Bergsbaken et al., 2009, Rathinam et al., 2012). This cell death pathway involves canonical (caspase-1) or non-canonical (caspase-11) activation of the inflammasome pathways (Rathinam et al., 2012, Bergsbaken et al., 2009). Activated caspase-1 or caspase-11 cleaves gasdermin D (GSDMD), which forms pores in the cell membrane enabling the release of intracellular contents, including pro-inflammatory cytokines. Pyroptosis is thought to play a crucial role in the clearance of bacterial and viral infections (Zhu et al., 2017). It has also been linked to the pathogenesis of chronic inflammatory conditions, such as colon cancer (Derangere et al., 2014), liver fibrosis (Wree et al., 2014), and atherosclerosis (Xu et al., 2018). In this context, IBD patients and animals subjected to experimental colitis showed increased epithelial GSDMD expression and genetic ablation of GSDMD attenuated colitis severity in mice (Bulek et al., 2020).

Here, we report that epithelial-specific deletion of *Gpr35* leads to reduced numbers of goblet cells in the proximal colon, which correlated with reduced *Muc2* expression. This resulted in microbiome alterations and increased susceptibility to the A/E pathogen *C. rodentium*. Mechanistically, epithelial *Gpr35* deficiency leads to activation of caspase-11-mediated pyroptosis in goblet cells. This study demonstrates that GPR35 is critical for the integrity of the colonic epithelial barrier.

## Results

### Epithelial *Gpr35* deletion reduces goblet cell numbers

To characterize the impact of Gpr35 on the epithelial barrier, we made use of the zebrafish (*Dario rerio*) model. Previously, we CRISPR-targeted two GPR35 paralogs in zebrafish, *gpr35a* and *gpr35b,* which revealed that the latter is more similar in function to human GPR35 (Kaya et al., 2020). Goblet cell numbers in *gpr35b* deficient (*gpr35b^uu19b2^*) zebrafish were decreased compared to Gpr35^wt^ larvae (Figures 1A and 1B). In contrast, Gpr35^wt^ larvae treated with the GPR35 agonists LPA and Zaprinast displayed increased goblet cell numbers indicated by Alcian blue staining, although the LPA treatment did not reach statistical significance (Figures 1C and 1D). To translate our findings from zebrafish into mice, we identified the GPR35-expressing cells in *Gpr35*-tdTomato reporter mice. Flow cytometry analysis revealed high GPR35 expression in colonic ECs (Figure S1A). Furthermore, *ex vivo* imaging of small and large intestine from *Gpr35*-tdTomato x *Cx3cr1*-GFP double reporter mice located GPR35 in CX3CR1^+^ lamina propria macrophages and epithelial cells (Figure S1B). Given the prominent expression of *Gpr35* in ECs, we set out to investigate whether epithelial *Gpr35* deficiency affects the epithelium. For this purpose, we crossed *Gpr35^f/f^* with *Villin1-Cre* mice to induce *Gpr35* deficiency in ECs (*Gpr35^f/f^Vil^+^*). The deletion of *Gpr35* from ECs in *Gpr35^f/f^Vil^+^* mice was confirmed by immunofluorescent staining for GPR35 (Figure S1C). PAS-Alcian blue staining revealed a decreased number of PAS^+^ goblet cells in the proximal colon of *Gpr35^f/f^Vil^+^* mice compared to *Gpr35^wt^* littermates (Figures 1E and 1F). Of note, this phenotype was absent in the distal colon (Figures S1D and S1E). Transmission electron microscopy analysis showed a high count of fused granules in mice lacking epithelial *Gpr35* indicating an abnormality in goblet cell appearance (Figures 1G and 1H). Immunohistochemistry (IHC) staining of Muc2 further demonstrated goblet cell depletion in *Gpr35^f/f^Vil^+^* mice (Figure 1I). Accordingly, *Muc2* mRNA expression levels in *Gpr35^f/f^Vil^+^* mice were significantly decreased compared to *Gpr35^wt^* littermate (Figure 1J). To investigate whether loss of epithelial *Gpr35* leads to further changes in colonic goblet cells that may not be detected in histological analysis, we measured the level of transcription factors involved in goblet cell differentiation. The expression of secretory lineage differentiation factor *Atoh1*, but not *Hes1*, was significantly reduced in *Gpr35^f/f^Vil^+^* mice compared to *Gpr35^wt^* littermates (Figure 1K). Furthermore, mRNA levels of the goblet cell maturation factors *Gfi1* and *Spdef* were downregulated in epithelial *Gpr35* deficient mice (Figure 1L).

**Figure 1.**
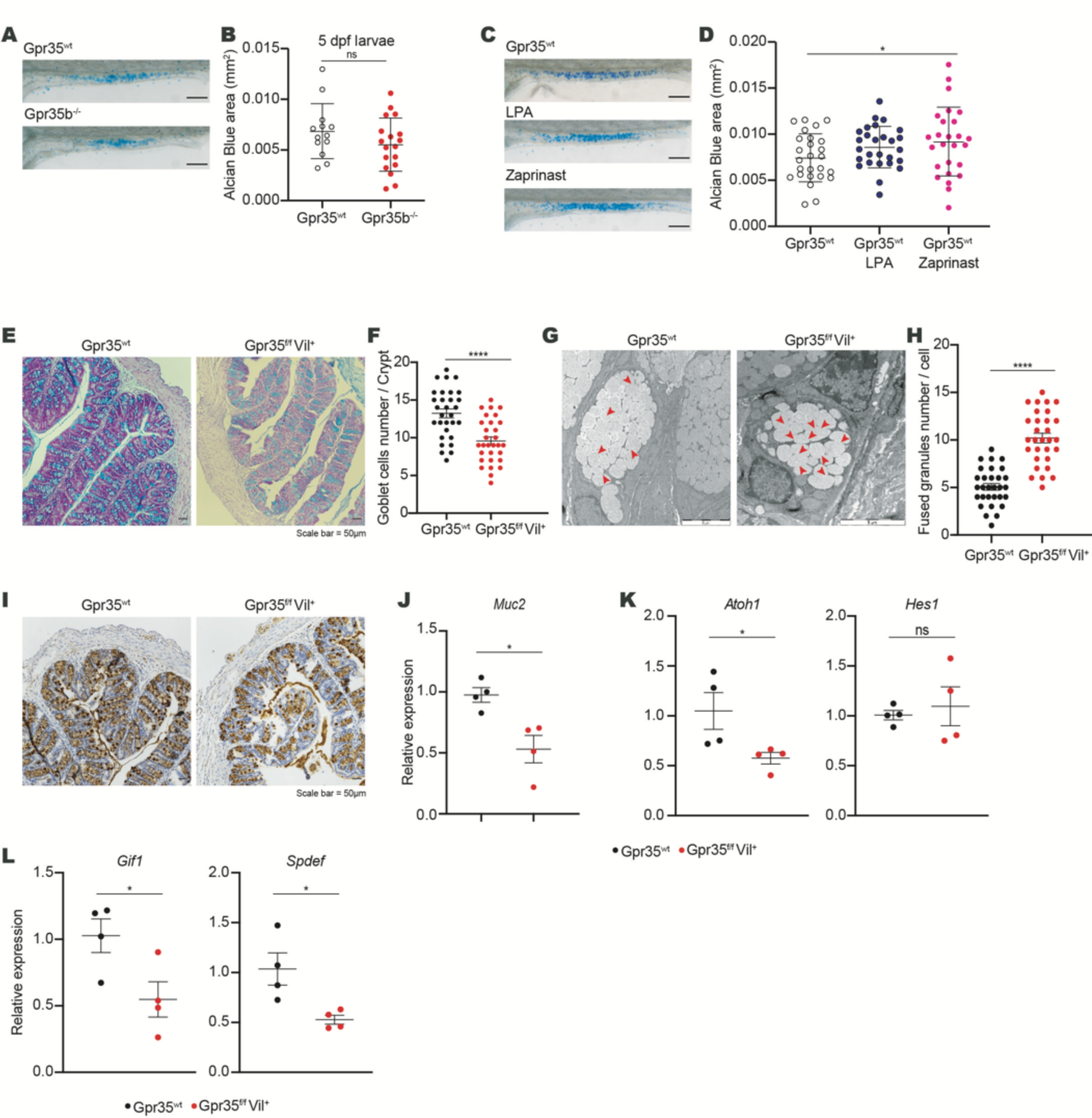
Epithelial cell-specific *Gpr35* deletion reduces goblet cell numbers. (A) Representative magnification of the intestine of either *Gpr35^wt^* or gpr35b-deficient zebrafish larvae showing Alcian blue staining. Scale bars, 100 µm. (B) Quantification of the Alcian blue area in (A) performed by using automatic Alcian blue color deconvolution in ImageJ. 1 dot = 1 larva. (C) Representative magnification of the intestine of *Gpr35^wt^* zebrafish larvae treated or not with either zaprinast or LPA and stained with Alcian blue. Scale bars, 100 µm. (D) Quantification of the Alcian blue area in (C) performed by color deconvolution. 1 dot = 1 larva. (E) Representative AB/PAS staining of proximal colon sections obtained from *Gpr35^f/f^Vil^+^* and *Gpr35^wt^* littermates. Scale bars, 50 µm. (F) Cell count of goblet cells in (E) performed blindly by two different investigators in at least 30 crypts. (G) Transmission electron microscopy analysis of goblet cell morphology in the proximal colon of *Gpr35^f/f^Vil^+^* and *Gpr35^wt^* littermates. Red arrowheads indicate fused granules. Scale bars, 5 µm. (H) Quantification of the number of mucin granules per goblet cell in (G) performed blindly by two different investigators in at least 30 cells. (I) Representative images of proximal colon sections obtained from *Gpr35^f/f^Vil^+^* and *Gpr35^wt^* littermates and stained for Muc2 protein by immunohistochemistry. Scale bars, 50 µm. (J-L) mRNA expression levels of (J) *Muc2*, (K) *Atoh1*, *Hes1*, (L) *Gif1* and *Spdef* measured by RT-qPCR in proximal colon samples obtained from *Gpr35^f/f^Vil^+^* (n = 4) and *Gpr35^wt^* littermates (n = 4). Each dot represents one animal with medians. Data are represented as mean ± SEM, ns not significant, *p ≤ 0.05, **p ≤ 0.01, ***p ≤ 0.001, ****p ≤ 0.0001 by Mann-Whitney U test.

### Epithelial cell-specific *Gpr35* deletion correlates with an altered mucus-associated microbiome

Given that an intact mucin barrier is essential for fencing off microorganisms from epithelial cells (Johansson et al., 2011b, Zarepour et al., 2013), we hypothesized that in *Gpr35^f/f^Vil^+^* mice, the microbiota would be in close proximity to epithelial cells in *Gpr35^f/f^Vil^+^* mice. Accordingly, 16S *in situ* hybridization (FISH) analysis of Carnoy fixed tissues revealed that the microbiota in these mice is in close contact with the epithelium (Figures 2A). This prompted us to investigate whether epithelial *Gpr35*-mediated goblet cell depletion impacts the microbial ecosystem. To compare the intestinal bacterial composition, we performed 16S rRNA amplicon sequencing on fecal as well as on mucosa-associated bacteria harvested from either *Gpr35^f/f^Vil^+^* animals or their *Gpr35^wt^* control littermates (see Workflow for details). We kept *Gpr35^f/f^Vil^+^* and *Gpr35^wt^* control mouse litters in separate cages after weaning since co-housing is a confounding variable for studying the gut microbiota. The mucosa-associated bacterial communities of *Gpr35^f/f^Vil^+^* were distinct from those of control littermates, as evidenced by principal coordinates analysis (PCoA) and the hierarchical clustering in young and aged mice (Figures 2B and S2A). Among these differences, *Deferribacteres* were overrepresented in *Gpr35^f/f^Vil^+^* mice versus *Gpr35^wt^* littermates (Figures 2C and S2B). Interestingly, *Clostridia* were more abundant in older control littermates than in *Gpr35^f/f^Vil^+^* mice (Figures 2C). At the genus level, *Mucispirillum*, a member of the *Deferribacteres* class, was enriched in the mucosa of *Gpr35^f/f^Vil^+^* mice compared to *Gpr35^wt^* littermates (Figure S2C). In addition, *Lachnospiraceae_NK4A136_group*, a member of the *Clostridia* class, was more abundant in *Gpr35^wt^* mice (Figure S2C). However, no differences were found between the fecal bacterial compositions of young *Gpr35^f/f^Vil^+^* and *Gpr35^wt^* mice (Figures 2D, 2E, S2D and S2E). In contrast, aging resulted in distinct rare taxa in *Gpr35^f/f^Vil^+^* mice compared to *Gpr35^wt^* littermates (Figures 2E and S2E). Taken together, these data suggest that reduction in goblet cells following epithelial *Gpr35* deletion is associated with changes in the mucosa-associated bacteria composition.

**Figure 2.**
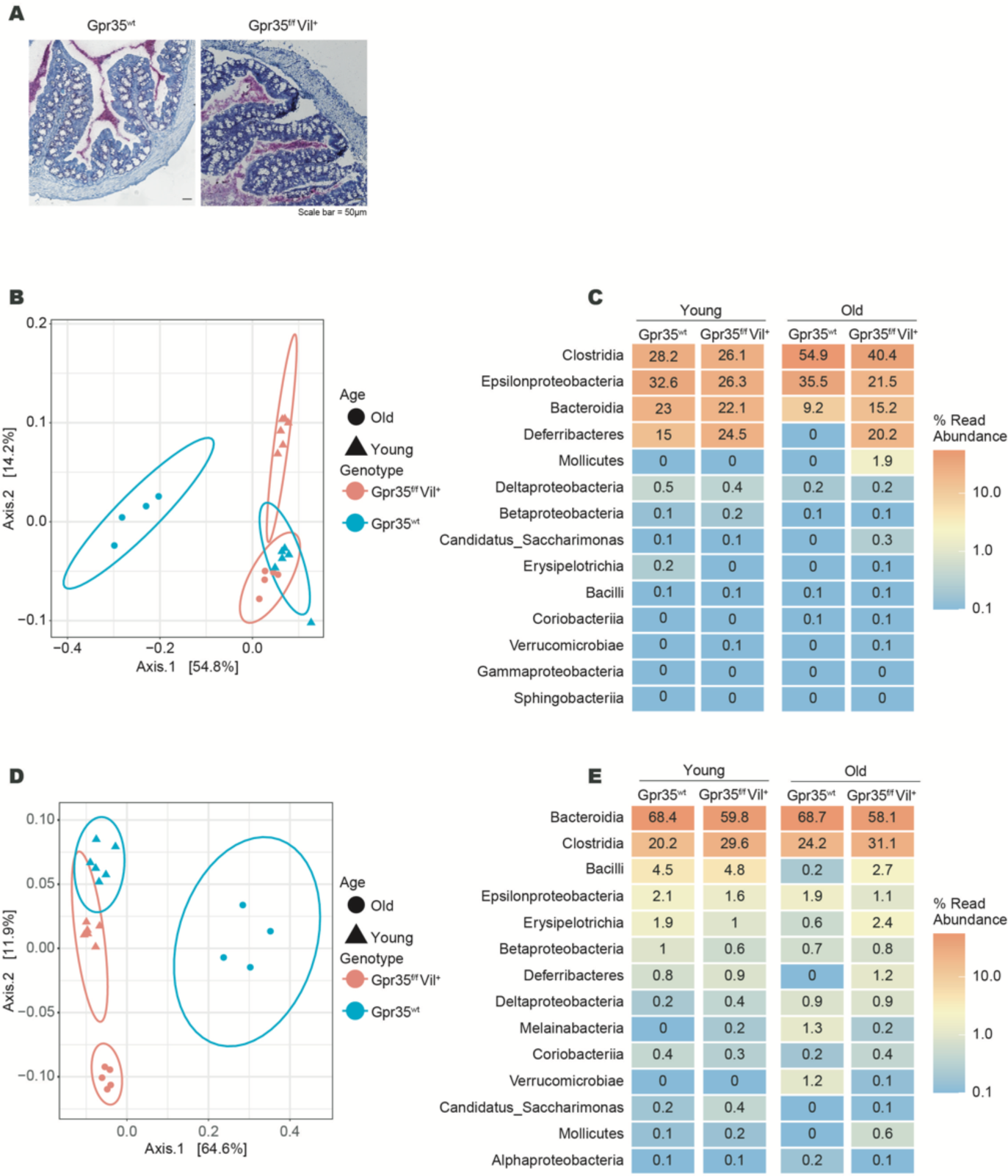
Epithelial cell-specific *Gpr35* deletion correlates with an altered mucosa-associated microbiome. (A) Visualization of bacteria in relation to the epithelium via 16S rRNA *in situ* hybridization (pink) in proximal colon sections obtained from *Gpr35^f/f^Vil^+^* and *Gpr35^wt^* littermates. Scale bars, 50 µm. (B) Principal component analysis based on Jaccard distance of the rarefied abundance of proximal colon mucosa-associated bacterial communities in *Gpr35^f/f^Vil^+^* young (n = 6), *Gpr35^f/f^Vil^+^* old (n = 5) and *Gpr35^wt^* young (n = 6), and *Gpr35^wt^* old (n = 4) littermates. (C) Relative abundance of taxonomic groups averaged across mucosa-associated bacteria samples of old and young *Gpr35^f/f^Vil^+^* and *Gpr35^wt^* littermates. (D) Principal component analysis based on Jaccard distance of rarefied abundance of fecal bacterial communities in *Gpr35^f/f^Vil^+^*young (n = 6), *Gpr35^f/f^Vil^+^* old (n = 5) and *Gpr35^wt^* young (n = 6), and *Gpr35^wt^* old (n = 4) littermates. (E) Relative abundance of taxonomic groups averaged across fecal samples of old and young *Gpr35^f/f^Vil^+^* and *Gpr35^wt^* littermates. Fecal and mucosal associated bacteria were collected from the same animals.

### Heterogeneity of colonic epithelial cells in *Gpr35^f/f^Vil^+^* mice

To gain further insight into the molecular mechanism by which deletion of *Gpr35* disrupts goblet cell function, we performed a single-cell RNA-sequencing (scRNA-seq) experiment using a 10X Genomics Chromium platform. We isolated proximal colon ECs from 4 *Gpr35^f/f^Vil^+^* and 4 *Gpr35^wt^* female littermates at steady-state, which were co-housed to minimize the effects of different microbiome compositions. We dissociated crypts into single-cell suspensions and sorted for CD326^+^, CD45^-^ and CD31^-^ cells (Figure S3A). After filtering out low-quality cells (see Methods), we retained 8,627 WT cells and 14,792 KO cells, ranging from 1,567 to 4,774 cells per sample (Figures S3B-S3D). Cells from both *Gpr35^f/f^Vil^+^* and *Gpr35^wt^* mice were combined and partitioned into 18 clusters (Figure 3A), which were annotated using whole-transcriptome comparison to reference scRNA-seq atlases of colonic and small intestinal epithelia (Haber et al., 2017, Parikh et al., 2019) (Figures 3B and 3C) and to reference the expression of known marker genes (Figures 3D-3I and S3E). Finally, the cells were assigned to 10 different cell types or subtypes, comprised of stem cells, transit-amplifying cells (separated into G1 and G2, according to their cell-cycle signature); (Figure S2F), absorptive colonocytes (with distinct stages of maturation: early than late progenitors, followed by immature and then mature colonocytes), goblet cells (immature, mature and crypt top cells), enteroendocrine and tuft cells. The annotation of the cells is shown on a *t*-distributed stochastic neighborhood embedding (*t*-SNE) (Figure 3J).

**Figure 3.**
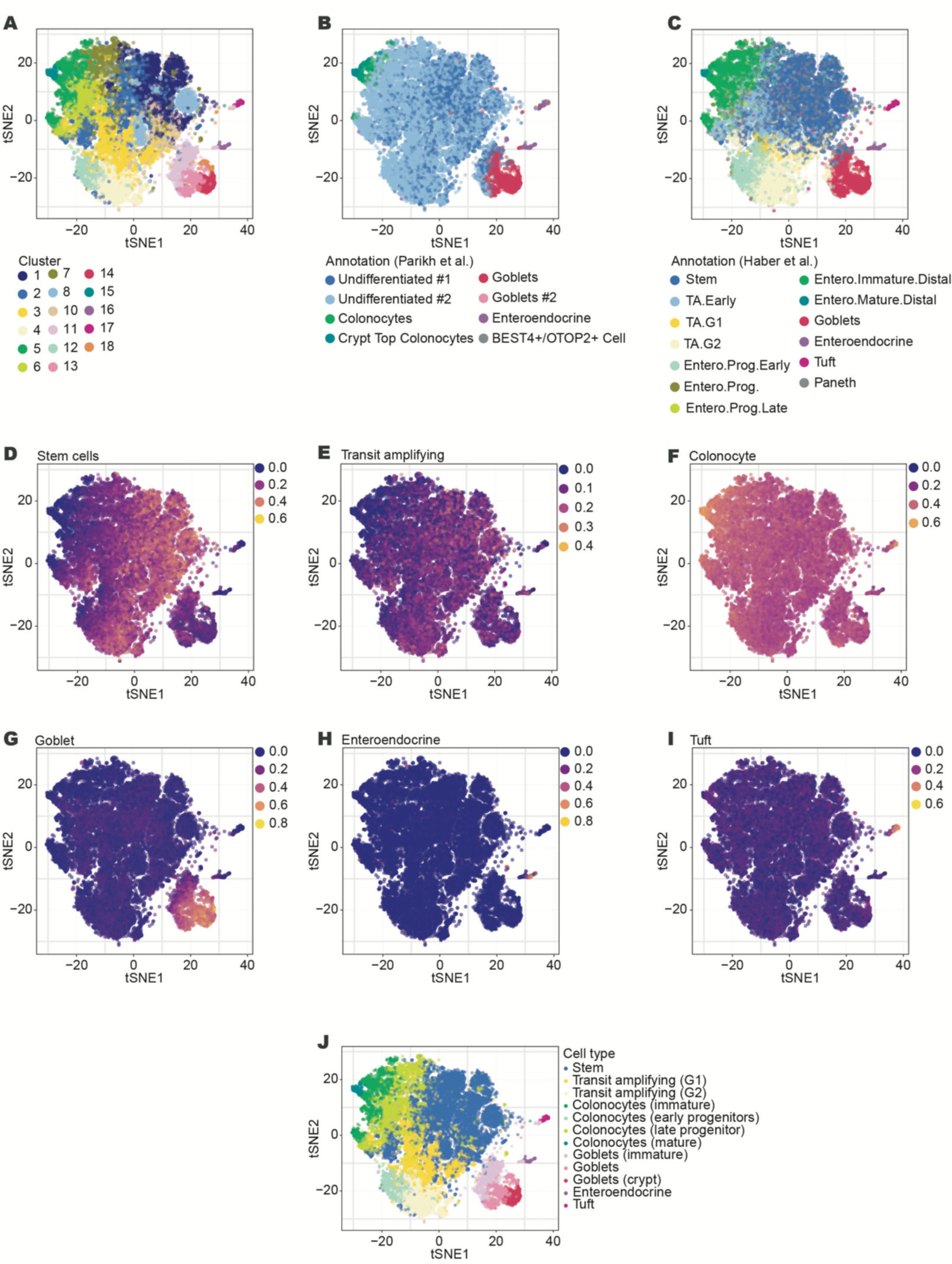
Heterogeneity of colonic epithelial cells in *Gpr35^f/f^Vil^+^* mice. (A-I) *t*-SNE plot showing proximal colonic epithelial cells from *Gpr35^f/f^Vil^+^*(n = 4) and *Gpr35^wt^* (n = 4) littermates assayed via scRNA-seq. (A) Partition of cells into hierarchical clusters. (B-C) Cell type annotation using whole-transcriptome comparison to reference scRNA-seq atlases of (B) colonic, (C) small intestinal epithelial cells. (D-I) Average expression score of known marker genes for different cell types. (J) Final cell-type annotation used in the paper.

### Increased pyroptotic signatures in goblet cells lacking epithelial *Gpr35*

We next turned to the analysis of differential gene expression between *Gpr35^f/f^Vil^+^* and *Gpr35^wt^* cells stratifying the analysis by cell type (see Methods). The genes differentially expressed in each cell type at a false discovery rate of 5% are shown in Table S1. Interestingly, goblet cells also showed the highest number of differentially expressed genes, with notably 14 genes significantly upregulated in *Gpr35^f/f^Vil^+^* compared to *Gpr35^wt^* cells (Figures 4A and S4A-S4C). A gene set enrichment analysis on Gene Ontology categories indicated an increased expression of genes related to pyroptosis in *Gpr35^f/f^Vil^+^* goblet cells (Figures 4B and 4C).

**Figure 4.**
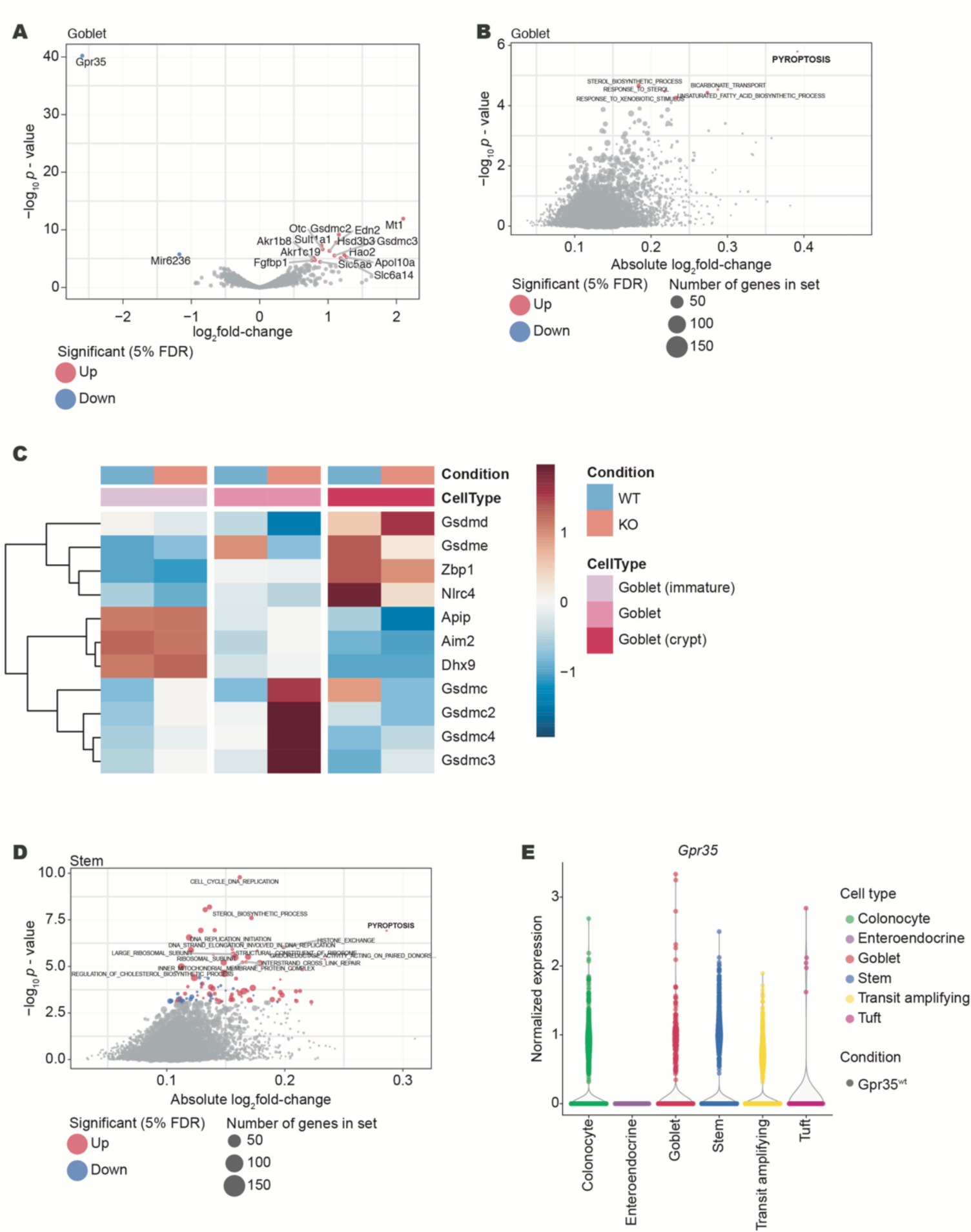
Increased pyroptotic signatures in goblet cells lacking epithelial *Gpr35*. (A) Volcano plot showing genes that are differentially expressed between *Gpr35^f/f^Vil^+^* (n = 4) and *Gpr35^wt^* goblet cells (n = 4). Grey dots highlight all genes analyzed; red or blue dots highlight genes significantly up or down- regulated genes in *Gpr35^f/f^Vil^+^* cells. (B) Gene set enrichment analysis on differential expression results from goblet cell clusters comparing *Gpr35^f/f^Vil^+^* to *Gpr35^wt^* cells. (C) Heatmap of genes from the Gene Ontology category pyroptosis, displaying the centered and scaled average expression across cells from the goblet cell clusters from both groups. (D) Gene set enrichment analysis on differential expression results from stem cell clusters comparing *Gpr35^f/f^Vil^+^* to *Gpr35^wt^* cells. (E) Normalized expression levels of Gpr35 in *Gpr35^wt^* cells from the scRNA-seq dataset, across annotated cell types.

This category was also upregulated in stem cells, although to a smaller extent (Figure 4D). We thus verified whether the differentiation rate of stem cells was affected. Colon length and crypt height was similar in *Gpr35^f/f^Vil^+^* and *Gpr35^wt^* littermates (Figure S4E-S4G). In addition, no differences were found in the mRNA expression levels of colonocyte (*Atpb1*, *Scla2*) and enteroendocrine (*Sct* and *Cck*) markers (Figure S4G). Furthermore, we found that the goblet cell cluster displays the highest *Gpr35* expression level among all clusters (Figure 4E). Overall, these results suggest that deletion of epithelial Gpr35 affects mostly goblet cells rather than other epithelial or intestinal stem cells.

### Pyroptosis upon *Gpr35* deletion is caspase-11 dependent

Pyroptosis is regulated by a canonical caspase-1-dependent or a non-canonical caspase-1- independent mechanism executed by caspase-11 (Rathinam et al., 2012). Upon activation, caspase-1 or caspase-11 directly cleaves GSDMD generating a 31-kDa N-terminal fragment, which initiates pyroptosis. To validate our scRNA-seq findings, we measured the protein level of GSDMD and found a significant increase in the cleaved form of both GSDMD and caspase- 11 but not caspase-1 in *Gpr35^f/f^Vil^+^* mice compared to *Gpr35^wt^* controls (Figures 5A, 5B and S5A-S5D). Consistent with these findings, *Gpr35^wt^* explants treated with a GPR35 inhibitor, ML194, had higher expression levels of cleaved GSDMD and cleaved caspase-11 (Figures 5C, 5D and S5E-5G). To further confirm that increase of pyroptosis signature is GPR35-dependent, we pretreated explants obtained from *Gpr35^wt^* mice with GPR35 agonists namely Zaprinast and LPA. Subsequently, explants were stimulated with *E. coli* outer membrane vesicles (OMVs), which have been described as vesicles secreted by Gram-negative bacteria that can induce pyroptosis via caspase-11 (Kaparakis-Liaskos and Ferrero, 2015, Russo et al., 2018). Immunoblotting analysis showed an increase in the expression level of both cleaved GSDMD and cleaved caspase-11 in OMV-treated *Gpr35^wt^* explant, which was rescued with Zaprinast but not LPA pre-treatment (Figures 5E, 5F and 5H-5J).

**Figure 5.**
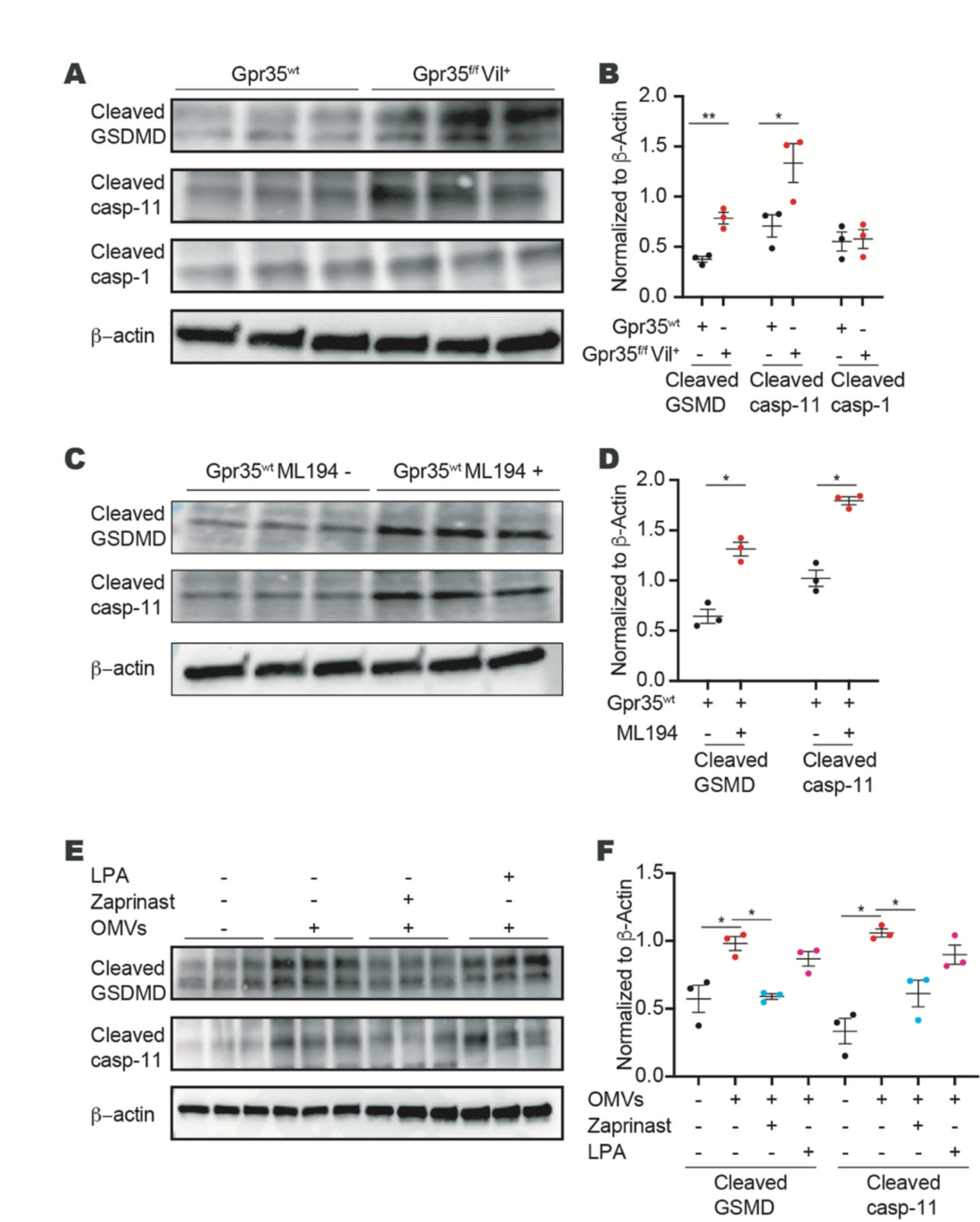
Pyroptosis upon *Gpr35* deletion is caspase-11 dependent. (A) Protein expression of GSDMD, cleaved caspase-11 and cleaved caspase-1 in crypt samples isolated from the proximal colon obtained from *Gpr35^f/f^Vil^+^* (n = 3) and *Gpr35^wt^* littermates (n = 3). (B) Densitometry analysis of (A). (C) Protein expression of GSDMD and cleaved caspase-11 in proximal colon explant obtained from *Gpr35^wt^* mice treated with ML194 at 10 µM for 3 h. (D) Densitometry analysis of (C). (E) Protein expression of GSDMD and cleaved caspase-11 in proximal colon explant obtained from *Gpr35^wt^* mice treated with either Zaprinast or LPA at 10 µM for 1.5 h followed by OMVs treatment at 10 µg for 1.5 h. (F) Densitometry analysis of (E). Each dot represents one animal with medians. Data are represented as mean ± SEM *p ≤ 0.05, **p ≤ 0.01, ***p ≤ 0.001, ****p ≤ 0.0001 by Mann-Whitney test.

### Epithelial GPR35 protects against *Citrobacter rodentium* infection

An intact mucus layer protects the host from the A/E pathogen *C. rodentium* (Bergstrom et al., 2010). Given that deletion of *Gpr35* in epithelial cells reduced goblet cell numbers, we hypothesized that Gpr35 would affect the course of *C. rodentium* infection. To test this, we subjected global *Gpr35* deficient mice (*Gpr35^-/-^*) and *Gpr35^wt^* mice to infection with *C. rodentium*. Although the *C. rodentium* kinetic clearance was comparable between *Gpr35^-/-^* and *Gpr35^wt^* animals, the bacterial counts were significantly higher in the feces of *Gpr35^-/-^* mice between days 3 and 9 post-infection (p.i.) (Figure S6A). *C. rodentium* load was lower in peripheral tissue of *Gpr35^wt^* mice at day 21 p.i. while higher dissemination to the mesenteric lymph node (MLN) and the liver was noted in *Gpr35^-/-^* mice (Figure S6B). To exclude the possibility that macrophages expressing GPR35 contribute to the clearance of *C. rodentium,* we investigated whether *Gpr35* deletion in macrophages versus ECs affects mice differently during infection with *C. rodentium*. To test this, we interbred floxed *Gpr35* locus mice (*Gpr35^f/f^*) with *Cx3cr1^CreER^* allowing a tamoxifen-inducible deletion of GPR35 (*Gpr35^ΔCx3cr1^*) in CX3CR1^+^ macrophages. Interestingly, unlike the phenotype observed in *Gpr35^-/-^* mice, the deletion of *Gpr35* in macrophages did not affect fecal bacterial load or bacterial dissemination to peripheral tissues (Figures S6C and S6D). Thus, GPR35 expressed by macrophages does not drive anti-*C. rodentium* defense, suggesting that the epithelial *Gpr35* deficiency might be responsible for the observed reduction in *C. rodentium* clearance. In contrast to *Gpr35^ΔCx3cr1^* mice, infected *Gpr35^f/f^Vil^+^* mice were characterized by a higher pathogen load in the stool (Figure 6A) and bacterial dissemination to the MLN or the liver (Figure 6B). Serum IgG levels were similar between mice lacking epithelial *Gpr35* and their control littermates (Figure 6C). In addition, *C. rodentium*-specific IgG levels in serum were similar between mice lacking epithelial *Gpr35* and their control littermates (Figure 6C). This suggested that the systemic adaptive immune response to *C. rodentium* was intact in *Gpr35^f/f^Vil^+^* mice and indicated an alternative mechanism.

**Figure 6.**
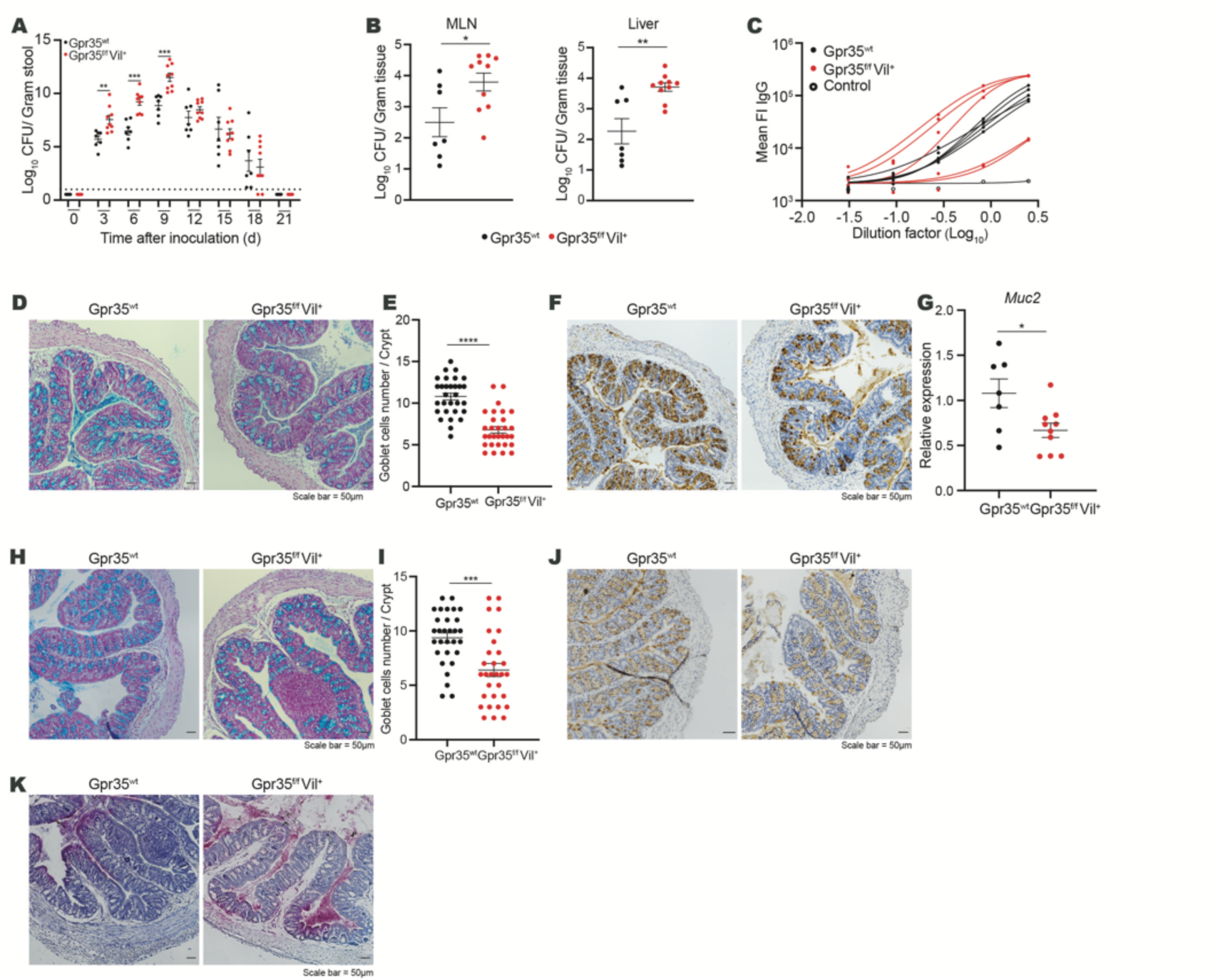
Epithelial GPR35 protects against *Citrobacter rodentium* infection. (A and B) *C. rodentium* CFU/g of (A) feces, (B) MLN and liver from *Gpr35^f/f^Vil^+^* (n = 10) and *Gpr35^wt^* littermates (n = 7). Mice were killed at day 21 p.i. (C) IgG serum level in *Gpr35^f/f^Vil^+^* and *Gpr35^wt^* littermates on day 21 p.i. (D) Representative AB/PAS staining of proximal colon sections obtained from *Gpr35^f/f^Vil^+^* and *Gpr35^wt^* littermates on day 21 p.i. Scale bars, 50 µm. (E) Cell count of goblet cells in (D) performed blindly by two different investigators in at least 30 crypts. (F) Representative images of proximal colon sections obtained from *Gpr35^f/f^Vil^+^* and *Gpr35^wt^* littermates on day 21 p.i and stained for Muc2 protein by immunohistochemistry. Scale bars, 50 µm. (G) mRNA expression levels of *Muc2* measured by qRT-PCR in proximal colon samples obtained from *Gpr35^f/f^Vil^+^* and *Gpr35^wt^* littermates on day 21 p.i. (H) Representative AB/PAS staining of proximal colon sections obtained from *Gpr35^f/f^Vil^+^* and *Gpr35^wt^* littermates infected with *C. rodentium*. Mice were killed at day 9 p.i. Scale bars, 50 µm. (I) Cell count of goblet cells in (H) performed blindly by two different investigators in at least 30 crypts. (J) Representative images of proximal colon sections obtained from *Gpr35^f/f^Vil^+^* and *Gpr35^wt^* littermates on day 9 p.i. and stained for Muc2 protein by immunohistochemistry. Scale bars, 50 µm. (K) Visualization of bacteria in relation to the epithelium via 16S rRNA fluorescence *in situ* hybridization (pink) in proximal colon sections obtained from *Gpr35^f/f^Vil^+^* and *Gpr35^wt^* littermates on day 9 p.i. Each dot represents one animal with medians. Data represent two independent experiments combined. Data are represented as mean ± SEM *p ≤ 0.05, **p ≤ 0.01, ***p ≤ 0.001, ****p ≤ 0.0001 by two-way ANOVA with Tukey’s multiple comparisons test in (A) or unpaired student’s t test in (B), (E), (G) and (I).

**Figure 7.**
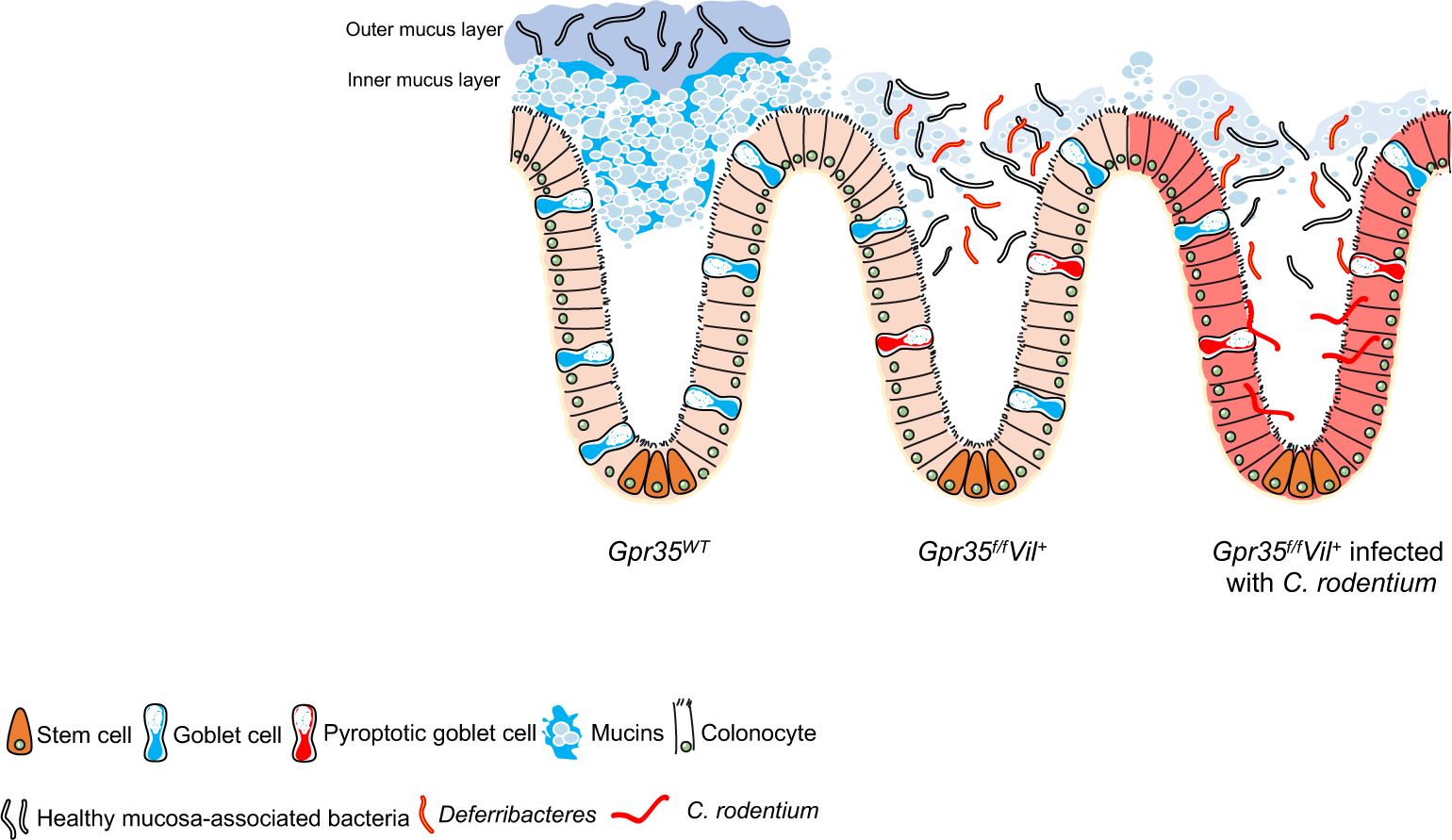
Goblet cells through their production and secretion of mucins play an essential role in protecting the host against luminal insults, and contribute to the mutualism between host and gut microbiota. Loss of epithelial *Gpr35* disrupts the goblet cell compartment by reducing their numbers and increasing pyroptosis levels. Dysregulation of goblet cells is correlated with changes in the mucosa-associated bacterial communities. As a result, *Gpr35^f/f^Vil^+^* mice displayed increased susceptibility to *C. rodentium-*induced colitis. (Adapted from SERVIER MEDICAL ART (CC of license 3.0)).

*C. rodentium* is an A/E pathogen that causes depletion of colonic goblet cells and their mucins (Bergstrom et al., 2008). Light microscopy analysis of PAS-Alcian blue-stained histological colon sections indicated decreased goblet cell numbers in the proximal colon of infected *Gpr35^f/f^Vil^+^* mice compared to WT littermates (Figures 6D and 6E). IHC analysis revealed fewer Muc2-positive cells (Figure 6F). Accordingly, *Gpr35^f/f^Vil^+^* mice displayed lower *Muc2* mRNA expression levels compared to *Gpr35^WT^* littermates (Figure 6G). Since the significant difference in pathogen load was observed on day 9 p.i., we measured the goblet cell number on this day. At this timepoint, goblet cell depletion in *Gpr35^f/f^Vil^+^* mice was more prominent, and *C. rodentium*-induced epithelial damage increased (Figures 6H and 6I). Lastly, IHC indicated fewer Muc2^+^ cells in mice lacking *Gpr35* in the epithelium (Figure 6J). 16S FISH analysis revealed the microbiota and *C. rodentium* were in closer proximity to the epithelium and confirmed more widespread infiltration of *C. rodentium* in *Gpr35^f/f^Vil^+^* mice compared to *Gpr35^wt^* littermates (Figures 6K). Collectively, these data indicate that the loss of *Gpr35* in ECs leads to an exacerbated bacterial burden associated with goblet cell depletion. These changes likely contribute to a barrier defect and lead to increased peripheral bacterial translocation upon *C. rodentium* infection.

Based on previous reports describing that goblet cells are critical in protection against DSS-induced colitis (Petersson et al., 2011), we finally sought to support our findings by exposing *Gpr35^f/f^Vil^+^* and *Gpr35^WT^* mice to DSS as a second experimental colitis model. Upon challenge with DSS, *Gpr35^f/f^Vil^+^* mice displayed more severe body weight loss (Figure S6E). There was no significant difference between *Gpr35^f/f^Vil^+^* and *Gpr35^WT^* mice in shortening of the large intestine (Figure S6F). The endoscopic evaluation indicated higher signs of colitis in *Gpr35^f/f^Vil^+^* compared to *GPR35^WT^* (Figure S6G). Consistently, histological analysis of H&E-stained colon tissue sections from *Gpr35^f/f^Vil^+^* mice showed significantly more mucosal damage, loss of goblet cells and inflammatory cell infiltrates than those from *Gpr35^WT^* mice (Figures S6H and S6I).

## Discussion

Goblet cells control host-microbe interaction through secretion of mucin. Impaired mucus production is associated with the development of UC. This study describes how the IBD risk gene GPR35 regulates mucosal barrier integrity through direct activity on goblet cells. We discovered that loss of epithelial *Gpr35* directly reduces goblet cell numbers and is associated with changes in the composition of mucosa-associated bacteria. We demonstrated that epithelial Gpr35 guards goblet cells from dysregulated pyroptosis, a cell death mode promoting inflammation (Bergsbaken et al., 2009).

Our data show that epithelial Gpr35 is essential for host defense against invasive bacterial infection. We observed that abrogated mucus production in mice lacking epithelial *Gpr35* results in higher susceptibility to the A/E pathogen *C. rodentium*. Accordingly, previous studies have demonstrated the critical role of goblet cells and mucus secretion in defending against bacterial pathogens of the A/E family, including *C. rodentium*, thereby reducing overall tissue damage (Bergstrom et al., 2010, Zarepour et al., 2013). Interestingly, susceptibility to *C. rodentium* increased at an early time point when the innate and not the adaptive immune system is in the process of clearing the infection, further indicating that impaired goblet cells contribute to the increased *C. rodentium* susceptibility in epithelial *Gpr35*-deficient mice. We have previously demonstrated that GPR35^+^ macrophages show higher transcript levels of pro-inflammatory cytokines, including *Il1b* and *Tnf* (Kaya et al., 2020). In turn, these cytokines have been shown to potentiate intestinal permeability (Neurath, 2014). These observations suggest that Gpr35 expressed by macrophages promote mucosal barrier loss and enhances bacterial invasion. Genetic deletion of *Gpr35* specifically in macrophages demonstrated that, unlike epithelial cells, Gpr35-expression in macrophages is not required for protection against *C. rodentium*.

Our results show an impairment of goblet cells in the proximal colon but not in the distal colon. A recent study elegantly showed that proximal colon-derived mucin governs the mucus barrier’s composition and function and is an essential element in regulating host-microbiota symbiosis through encapsulating microbiota-containing fecal pellets by a mucus layer mainly derived from the proximal colon (Bergstrom et al., 2020). *Gpr35^f/f^Vil^+^* mice were more susceptible to DSS-induced colitis, a model that appears to be more severe in the distal (Randhawa et al., 2014) colon. Possibly, O-glycan-rich mucus derived from goblet cells of the proximal colon may influence DSS induced-colitis in the distal colon, and thus a reduction of goblet cells in the proximal colon might explain the exacerbated DSS colitis observed in Gpr*35^f/f^Vil^+^* mice.

Previous studies have indicated that GPR35 promotes the mucosal barrier by using either agonist/antagonist (Tsukahara et al., 2017) or GPR35 global knockout mice (Farooq et al., 2018). However, these studies lacked the genetic models required to dissect both the cell-specific protective effects of GPR35 and the underlying mechanisms of these effects. We analyzed mice during steady-state to explore whether epithelial GPR35 is the main factor affecting the epithelial integrity since severe inflammation would result in epithelial cell depletion. Interestingly, *Gpr35^f/f^Vil^+^* mice were characterized by decreased functional goblet cell count. Furthermore, the loss of epithelial *Gpr35* leads to a reduced expression of the downstream effectors *Gfi1* and *Spdef*, which are constitutively expressed in mature goblet cells and are crucial factors for goblet cell maturation (Nowarski et al., 2015).

Our findings indicate that *Muc2* was downregulated by loss of epithelial *Gpr35* leading to a thinner mucus layer and a closer microbe-epithelium interaction. It has been reported that deletion of *Muc2* leads to an imbalance of fecal bacterial composition in mice of different ages, mainly marked by increased Firmicutes and lower abundance of *Lachnospiraceae* (Wu et al., 2018). In agreement with this study, our 16S RNA sequencing analysis indicates that epithelial loss of *Gpr35* leads to fecal bacterial changes in old mice. Taxa of the *Deferribacteres* phylum, particularly *Mucispirillum Schaedleri,* were abundant in the mucosa of both young and old *Gpr35^f/f^Vil^+^* mice. Interestingly, this bacterium protects *Agr2^-/-^* mice against colitis by conferring resistance against *Salmonella* but not against *C. rodentium* induced infection (Herp et al., 2019). In line with our findings, the combined loss of the two IBD risk genes, NOD2 and the phagocyte NADPH oxidase CYBB, led to a selectively higher presence of *Mucispirillum Schaedleri* caused by an impairment of both neutrophil recruitment and NADPH oxidase activity (Caruso et al., 2019). Despite the reduced goblet cell numbers and *Muc2* expression in the proximal colon and increased *Mucispirillum Schaedleri* abundance in mice lacking epithelial *Gpr35*, we did not observe any signs of spontaneous colitis in these animals. Therefore, it is tempting to hypothesize that NADPH oxidase may compensate for Gpr35- mediated goblet cell depletion. Thus, whether double deficient *Gpr35^f/f^Vil^+^/Cybb*^-/-^ mice develop spontaneous colitis merits further investigation.

Our scRNA-seq showed that goblet cells express the highest level of *Gpr35* among all epithelial cell types, and thus the deletion of epithelial *Gpr35* affects mainly these cells. Furthermore, gene set enrichment analysis indicated an upregulation of pyroptosis genes, particularly in goblet cells. We found an elevated level of cleaved GSDMD in goblet cells of *Gpr35^f/f^Vil^+^* mice. Accordingly, GSDMD is upregulated in the colon of colitis mice and mucosal biopsies from IBD patients (Bulek et al., 2020). The same study showed that lack of GSDMD effectively reduces the severity of DSS-induced colitis (Bulek et al., 2020). In our study, we found that pyroptosis activation in goblet cells is caspase-11 dependent. A recent report showed that inhibition of caspase-11 reduces enteric infection-induced neuronal loss (Matheis et al., 2020).

The activation of GPR35 by Zaprinast inhibited pyroptosis induced by *E. coli* OMVs. These findings highlight the possibility that in the absence of epithelial *Gpr35* and, upon bacterial invasion, GSDMD mediates goblet cell pyroptosis to remove pathogen-infected cells. Consistently, it has been reported that intestinal epithelial cells can physically expel themselves from the epithelium via pyroptosis to prevent intracellular pathogens from breaching the epithelial barrier (Rauch et al., 2017, Zhu et al., 2017). Whether the increased abundance of *Mucispirillum Schaedleri* in *Gpr35^f/f^Vil^+^* mice is related to pyroptotic goblet cells requires further investigation.

In conclusion, we propose that epithelial Gpr35 maintains the barrier integrity by preserving goblet cells which are indispensable for the defense against intestinal pathogens. This work suggests that pharmacological modulation of Gpr35 signaling may represent a novel strategy to prevent the breakdown of epithelial barrier integrity. Understanding the relationship between the microbiota and goblet cells might provide promising strategies to treat IBD patients suffering from dysregulated epithelial barrier integrity.

## Material and Methods

### Experimental models

Zebrafish WT and gpr35b mutants (Kaya et al., 2020), in AB genetic background, were kept at the Karolinska Institute Zebrafish Core Facility, Sweden. Breeding and experiments were performed under ethical permits Nr 5756/17 and Nr 14049/19, conferred by the Swedish Board of Agriculture (Jordburksverket).

### Transgenic animal models

*Gpr35*-tdTomato, *Gpr35*^−/−^ and *Gpr35^f/f^* animals were constructed as previously described (Kaya et al., 2020). *Gpr35*-tdTomato mice were crossed with *Cx3cr1*-GFP mice to generate the double reporter mouse line. *Gpr35^f/f^* mice were crossed with Cre expressing lines: *Villin1 Cre (B6.Cg-Tg(Vil1-cre)^997Gum/J^*; (kindly provided by Claudia Cavelti, Department of Biomedicine, University of Basel) to target intestinal epithelial cells and *Cx3cr1^CreER^* (B6.129P2(Cg)-*Cx3cr1^tm2.1(cre/ERT2)Litt^*^/WganJ^) to target Cx3cr1^+^ lamina propria macrophages under a tamoxifen-inducible system. Genotyping was performed according to the protocols established for the respective strains. All strains were maintained at the animal facility of the Department of Biomedicine, University of Basel, Switzerland, and kept under specific pathogen-free conditions. Mice were fed a standard chow diet, and only females between 7- 12 weeks of age were selected for experimental groups. Animal experimentation was conducted under the Swiss Federal and Cantonal regulations (animal protocol number 3000 (canton Basel-Stadt)).

### Method details Tamoxifen treatment

*Gpr35^ΔCx3cr1^* and their respective littermates *Gpr35^wt^* were administered 75 mg tamoxifen (MedChemExpress #HY-13757A/CS-2870) / kg body weight dissolved in corn oil (Sigma- Aldrich #C8267) via intraperitoneal injection (i.p.) daily from day three before infection until day six post-infection, then injections were repeated every third day.

### *Ex vivo* imaging of colonic tissues

Colon was flushed with phosphate-buffered saline (PBS) (Sigma-Aldrich #D8537), opened longitudinally, and placed on a slide. A drop of PBS was added to prevent the tissue from drying, and the tissue was covered with a coverslip. Slides were imaged on a Nikon A1R confocal microscope.

### In vivo C. rodentium infection

RF-*C. rodentium* was generated as previously published (Manta et al., 2013). Prior to infection, bacteria were propagated overnight in LB broth medium supplemented with 300 µg/ml erythromycin. The next day, bacteria were pelleted, washed, and resuspended in PBS. Female mice (7-12 weeks) were gavaged with 2 x 10^9^ colony-forming units (CFU) of RF-*C. rodentium*. To calculate bacterial CFU from the feces, liver, and MLN, samples were collected, homogenized in 1 ml PBS and clarified at 50xg for 1 min. Bacteria-containing supernatants were serially diluted, spotted in triplicate on LB agar erythromycin plates, and incubated at 37°C in a humidified atmosphere for 18 h. CFU counts were normalized to the weight of the sample.

### *C. rodentium*-specific IgG levels measurement

*C. rodentium* were cultured overnight as described above. Cultured bacteria were washed with PBS containing 2% BSA and 0.02% azide and incubated with diluted serum for 30 min. After washing twice in PBS containing 2% BSA, the bacteria were pelleted and stained with mouse anti IgG and Pyronin Y for 30 min at 4°C. Bacteria were then pelleted down and fixed with 2% PFA. Immunoglobulin bound bacteria were analyzed by CytoFLEX (Beckman coulter). **Experimental colitis** For acute experimental colitis, weight-matched 7 to 12-week-old female mice were administered with 2% DSS (M.W. 36,000-50,000 Da; MP Biomedicals #160110) in their drinking water ad libitum for five days followed by two days of normal drinking water.

### Mouse Endoscopy

To assess macroscopic colitis severity, mice were anaesthetized with 100 mg/kg bodyweight ketamine and 8 mg/kg bodyweight Xylazine intraperitoneally. The distal 3 cm of the colon and the rectum were examined with a Karl Storz Tele Pack Pal 20043020 (Karl Storz Endoskope, Tuttlingen, Germany) as previously described (Melhem et al., 2017).

### Histology

After sacrifice, proximal and distal colon were removed, cleaned with PBS, fixed with 4% paraformaldehyde (PFA) (Sigma-Aldrich #F8775), and embedded in paraffin. Five µm-thick sections were stained with Hematoxylin-eosin. Histological scores for colonic inflammation were assessed semi-quantitatively using the following criteria (Kaya et al., 2020): mucosal architecture (0: normal, 1-3: mild-extensive damage); cellular infiltration (0: normal, 1-3: mild- transmural); goblet cell depletion (0: no, 1: yes); crypt abscesses (0: no, 1: yes); extent of muscle thickening (0: normal, 1-3: mild-extensive). To preserve goblet cells and mucus layer, colon biopsies were directly submerged in Carnoy’s fixative (60% Methanol, 30% Chloroform and 10% acetic acid) at 4°C overnight. Fixed tissues were embedded in paraffin and 5 µm- thick sections were stained with Alcian blue/PAS (Sigma-Aldrich #B8438). Images were acquired with Nikon Ti2 inverted microscope, and data were analyzed using FIJI software. Histological score and goblet cell count were assessed blindly by at least two investigators.

### Immunohistochemistry and immunofluorescence staining

Tissues from the colon’s proximal part were washed, fixed in 4% PFA, paraffin-embedded, and cut into 5 µm sections. Slides were deparaffinized in xylene, rehydrated in graded alcohols, and incubated in citrate buffer solution (pH = 6) for 20 min in a pressure cooker for antigen retrieval. Endogenous peroxidases were blocked with 3% hydrogen peroxide (Roth #9681.4) for 10 min at room temperature followed by 1 h blocking step with PBS containing 0.4% Triton X-100 5% goat serum (all Sigma-Aldrich) before incubation overnight at 4°C with anti-Muc2 (Novus Biologicals #NBP1-31231, 1:1000). The next day, slides were washed and incubated for 1 h at room temperature with anti-rabbit horseradish peroxidase-conjugated antibodies (Jackson ImmunoResearch #111-035-003, 1:500). Peroxidase activity was detected using 3,3’ Diaminobenzidine substrate (BD Pharmingen #550880). Slides were counterstained with hematoxylin, dehydrated, and mounted. Images were acquired with a Nikon Eclipse Ti2 microscope, and data were analyzed using FIJI software.

For immunofluorescence, slides were treated as described above except for the blocking of endogenous peroxidases step. Tissue sections were stained with rabbit polyclonal anti-mouse Gpr35 primary antibody overnight and goat anti-rabbit IgG secondary antibody. Slides were washed and then incubated for 1 h with Alexa Fluor 647 donkey goat anti-rabbit IgG (Life technologies #A21244, 1:500). NucBlue™ Live Cell Stain (Thermo Fisher #R37605) was used for nuclear staining, and samples were imaged using a Nikon A1R confocal microscope. Brightness and contrast settings were maintained between control and test images using NIS software.

### 16S fluorescence *in situ* hybridization

16S rRNA FISH was performed using the view RNA tissue assay core kit (Thermo Fisher #19931) according to the manufacturer’s instructions. Briefly, mice were sacrificed, and the proximal colon was harvested, cleaned from excess fat and feces, and placed in ice-cold Carnoy’s solution at 4°C overnight. The tissue was then washed twice in 100% ethanol for 15 min, followed by two washing steps in xylenes for 15 min, then embedded in paraffin and cut to 5 µm. Tissue sections were deparaffinized by heating to 65°C for 1 h followed by 5 min incubations with xylene (3x) then 100% ethanol (2x) at room temperature. Excess of ethanol was removed by incubating the slides at 40°C for 5 min. Slides were next heated in pretreatment solution for 15 min at 90°C and then washed in ddH2O (2x) for 1 min. Following this, slides were exposed to protease digestion (1:100) for 15 min at 40°C, washed with PBS (2x), fixed with 4% PFA for 3 min at room temperature, and washed in PBS to eliminate the excess of PFA. Next, slides were incubated for 2.5 h at 40°C with a bacterial 16S-DNA probe (Thermo Fisher #VX-01-14303) diluted to 1:40 in a prewarmed probe set dilution QT solution. After 3x washing steps, slides were exposed sequentially to the preamplifier hybridization, amplifier hybridization, and label probe 1-AP hybridization solutions for 40 min at 40°C with washing steps 2 min after each incubation. Following these steps, slides were incubated with the fast-red substrate for 30 min at 40°C, washed in PBS (1x), fixed in 4% PFA, and then washed again in PBS (1x). Slides were counterstained with hematoxylin, dehydrated, rinsed with deionized water, and incubated with NucBlue™ Live Cell Stain (Thermo Fisher #R37605) before mounting of coverslips. Images were acquired with Nikon A1R confocal microscope, and data were analyzed using FIJI software.

### RNA extraction and quantitative PCR

Mice were sacrificed, and the proximal colon was dissected. According to the manufacturer’s instructions, RNA was extracted from tissue using TRI Reagent (Zymo Research #R2050-1- 200) or RNeasy mini kit (Qiagen #74104). Genomic DNA was eliminated with RNase-Free DNase Set (Qiagen #79254), and 1g total RNA was reverse-transcribed and amplified using High Capacity cDNA Reverse Transcription (Applied Biosystems) kit. Quantitative PCR was performed using primers listed in Table S2 and TakyonLow Rox SYBR MasterMix blue (Eurogentec #UF-LSMT-B0701). Samples were run on an ABI ViiA 7 cycler. Amplifications were performed in duplicate, and Ct values were normalized to *Gapdh.* Relative expression was calculated by the formula 2^(-ΔCt).

### Colonic epithelial cells isolation, flow cytometry, and cell sorting

Mice were sacrificed, and the proximal colon was extracted and cleaned from excess fat and feces. The tissue was cut into 5 mm fragments and washed with ice-cold PBS until the supernatant became clear. Following incubation for 15 min in 5 mM EDTA-PBS solution, at 37°C, colonic crypts were released by shaking 15 times in ice-cold PBS. Crypts were further digested in Roswell Park Memorial Institute (RPMI) 1640 (Sigma-Aldrich) containing 0.5 mg/ml Collagenase type VIII (Sigma-Aldrich # R8758) and 10 U/mL DNase (Roche #04536282001) for 15 min at 37°C in a shaking water bath with 30 s vortexing each 5 min. The cell suspensions were filtered through a 70 µm cell strainer (Sarstedt #83.3945.070) and incubated for 30 min at 4°C with fixable viability dye eFluor455UV (eBioscience #65-0868) for live/dead cell exclusion. Cells were washed in PBS containing 2% Fecal Bovine Serum (FBS), 0.1% sodium azide, 10 mM EDTA (FACS buffer), and stained for surface antigens for 20 min at 4°C. Flow cytometric analysis was performed on a Fortessa flow cytometer (BD Biosciences). Epithelial cell staining was performed using Epcam^+^ (BioLegend #118213) and CD45^-^ (eBioscience #64-0451-82).

For single-cell RNA-sequencing, single-cell suspensions from four WT and *Gpr35^f/f^Vil^+^* littermate mice were stained with Epcam^+^ (BioLegend #118213), CD45^-^ (eBioscience #64- 0451-82) and CD31^-^ (BioLegend #102414) and sorted into Eppendorf tubes containing 50 µL of 1X PBS with 0.4% BSA and 5% FBS.

### Single-cell RNA sequencing

After sorting viable CD321^+^ CD45^-^ and CD31 cells from *Gpr35^f/f^Vil^+^* and *Gpr35^wt^* littermate mice, cells suspension volumes with a targeted recovery of 10,000 cells were loaded on 8 wells of a single 10X Genomics Chromium Single Cell Controller (one well per replicate), and 3’end libraries were generated using v3 chemistry. Libraries were sequenced on a flow-cell of an Illumina NovaSeq 6000 sequencer at the Genomics Facility Basel of the ETH Zurich (with 91nt-long R2 reads).

Data analysis was performed by the Bioinformatics Core Facility, Department of Biomedicine, University of Basel, Switzerland. Read quality was assessed with the FastQC tool (version 0.11.5). Sample and cell demultiplexing, read pseudo-alignment to the mouse transcriptome (Ensembl release 97)(Yates et al., 2020), and generation of the table of UMI counts were performed using Kallisto (version 0.46.0) and BUStools (version 0.39.2) (Melsted et al., 2019b, Melsted et al., 2019a). Further processing of the UMI counts table was performed by using R 4.0 and Bioconductor 3.11 packages, notably DropletUtils (version 1.8.0) (Griffiths et al., 2018, Lun et al., 2019), scran (version 1.16.0), and scater (version 1.16.2) (McCarthy et al., 2017), following mostly the steps illustrated in the Bioconductor OSCA book (https://osca.bioconductor.org/) (Lun et al., 2016, Amezquita et al., 2019). Based on the observed distributions, cells with 0% or more than 10% of UMI counts attributed to the mitochondrial genes (Ilicic et al., 2016), with less than 1,000 UMI counts, or with less than 631 detected genes were excluded. A total of 15,785 KO cells (ranging from 2,243 to 5,029 cells per sample) and 9,070 WT cells (ranging from 1,637 to 2,645 cells per sample) were used in the next steps of the analysis. Low-abundance genes with less than 0.01 UMI count on average across cells were excluded (11,994 genes were used in the next steps of the analysis). UMI counts were normalized with size factors estimated from pools of cells created with the scran package *quickCluster()* function (Lun et al., 2016, Vallejos et al., 2017). To distinguish between genuine biological variability and technical noise, we modeled the log-expression variance across genes using a Poisson-based mean-variance trend. The scran package *denoisePCA()* function was used to denoise log-expression data by removing principal components corresponding to technical noise. A *t*-distributed stochastic neighbor embedding (t-SNE) was built with a perplexity of 100 using the top 500 most variable genes and the denoised principal components as input. The cell cycle phase was assigned to each cell using the scran package *cyclone* function and the available pre-trained set of marker pairs for the mouse (Scialdone et al., 2015). Clustering of cells was performed using hierarchical clustering on the Euclidean distances between cells (with Ward’s criterion to minimize the total variance within each cluster; package cluster version 2.1.0). Cell clusters were identified by applying a dynamic tree cut (package dynamicTreeCut, version 1.63-1), which resulted in 18 clusters. The scran package *findMarkers()* function was used to identify marker genes up-regulated in each cluster or annotated cell type. The potential presence of doublet cells was investigated with the scran package *doubletCluster()* function and with the scDblFinder package (version 1.2.0) (Xi and Li, 2020). Consistent with the moderate loading of 10X wells, a low number of potential doublets was detected, and no further filtering was performed. The package SingleR (version 1.2.4) was used for reference-based annotation of cells (Aran et al., 2019). As a reference, we first used a scRNA-seq dataset from mouse small intestinal epithelial cells (GEO accession GSE92332) (Haber et al., 2017) aggregated into pseudo-bulk samples based on the provided cell-type annotation and transformed to log2CPM (counts per million reads) values. Secondly, a scRNA-seq dataset from human colon epithelial cells was used (GEO accession GSE116222) (Parikh et al., 2019). UMI counts from single cells were aggregated into pseudo-bulk samples based on the cell-type annotation and patient status (healthy, UC non-inflamed, or UC inflamed) obtained by personal correspondence to the authors and transformed to log2CPM (counts per million reads) values. The correspondence to mouse gene IDs was made by retrieving the 1-to-1 orthologs to human genes in the original dataset from Ensembl Compara (Herrero et al., 2016). SingleR pruned labels were retained to annotate cells to these pseudo- bulk reference datasets (Figures 3B and 3C), and a consensus annotation was manually derived from these results (Figure 3J). Complementary to this approach, a marker-based approach was used for annotation, where the averaged scaled expression of known markers was visualized on the t-SNE (Figures 3D-3I): Lgr5, Ascl2, Axin2, Olfm4 and Slc12a2 for stem cells, Bmi1, Lrig1, Hopx and Tert for transit-amplifying (TA) cells, Epcam, Krt8, Vil1, Alpi, Apoa1, Apoa4 and Fabp1 for enterocytes, Muc2, Clca1, Tff3 and Agr2 for goblet cells, Chga, Chgb, Tac1, Tph1 and Neurog3 for enteroendocrine cells, and Dclk1, Trpm5, Gfi1b, Il25, Klf3, Gng13 and Rgs2 for tuft cells. Based on the observation that cluster 9 displayed no precise specific marker gene, a relatively high expression of mitochondrial genes, and its cells were spread on the reduced dimension embeddings, we concluded that it was composed of low-quality cells and excluded it from further analyses (1,436 cells).

Differential expression between KO and WT cells stratified by annotated cell type was performed using a pseudo-bulk approach, summing the UMI counts of cells from each cell type in each sample when at least 20 cells could be aggregated. For goblet cells and colonocytes, progenitors, immature and mature cells subtypes were grouped to get sufficient cell numbers. Similarly, transit-amplifying G1 and G2 were grouped. Enteroendocrine and tuft cells could not be tested due to an insufficient number of cells. The aggregated samples were then treated as bulk RNA-seq samples (Lun and Marioni, 2017) and for each pairwise comparison, genes were filtered to keep genes detected in at least 5% of the cells aggregated. The package edgeR (version 3.30.3) (Robinson et al., 2010) was used to perform TMM normalization (Robinson and Oshlack, 2010) and to test for differential expression with the Generalized Linear Model (GLM) framework. Genes with a false discovery rate (FDR) lower than 5% were considered differentially expressed. Gene set enrichment analysis was performed with the function camera (Wu and Smyth, 2012) on gene sets from the Molecular Signature Database collections (MSigDB, version 7.2) (Liberzon et al., 2015, Subramanian et al., 2005). We considered only sets containing more than 10 genes, and gene sets with an FDR lower than 5% were considered significant.

The scRNA-seq dataset isavailable on the GEO repository under accession GSE169183.

### Immunoblotting

Following colonic crypts homogenization, as described above, total protein was extracted by lysing tissue in ice-cold RIPA buffer supplemented with protease inhibitor cocktail (Santa Cruz #sc-24948), sodium orthovanadate, and PMSF. Protein concentrations were quantified using the BCA method. For each group, 15 µg of protein were transferred to a nitrocellulose membrane after electrophoretic separation. The membranes were blocked using 5% of either dry milk or BSA in Tris Buffered Saline + Tween20 (TBS-T) buffer. The nitrocellulose membrane was then incubated overnight with the following primary antibodies: cleaved GSDMD (Cell Signaling Technology, #50928), cleaved caspase-11 (Abcam, #ab180673), cleaved caspase-1 (Invitrogen #AB 5B10), and β-actin (BD Biosciences #612656) at 1:1000 dilution. After washing steps in TBST, the membrane was incubated at room temperature for 1 hour with anti-rabbit IgG (H+L) or anti-mouse IgG (H+L) (both Jackson ImmunoResearch) at 1:15000 dilution. Proteins were visualized using SuperSignalTM West Femto or SuperSignal West Pico PLUS (both Thermo Fisher) chemiluminescent detection kits.

### 16S RNA sequencing

Fecal pellets were collected, weighed, and stored in cryo-storage vials at -80°C until processing. For mucosal-associated bacteria isolation, the proximal colon was dissected, opened longitudinally, and washed in PBS until no fecal matter was observed. The mucosal layer was manually scraped, weighed, and agitated for 20 minutes at 3000 rpm in 1mM ice-cold DL-dithiothreitol (DTT). After discarding the undissolved tissue, supernatants were centrifuged at 10,000g for 3 minutes, and sediments were collected. According to the manufacturer’s instructions, bacterial genomic DNA was isolated from sediments or fecal pellets using a DNeasy PowerSoil Kit (Qiagen #12888-100). Bacterial genomic DNA was used as a PCR template using previously established primers for the 16S rRNA gene (Table S3). PCR was performed in 25 µl reaction mix containing 2x KAPA HIFI HotStart Ready-mix (Roche #07958935001), 10µM primers and 10 ng of gDNA template from stool or tissue under the following conditions: 94°C for 3 minutes, followed by 27 cycles (for mucosal-associated bacteria samples) or 18 cycles (for feces samples) of 94°C for 30 seconds; 53°C (mucosal-associated bacteria samples) or 59°C (feces samples) for 15 seconds and 72°C for 15 seconds; after which a final elongation step at 72°C for 5 minutes was performed. After amplification, PCR products were cleaned with the Agencourt AMPure XP kit (Beckman Coulter #A63881), and amplicons were used to perform the index PCR using the Nextera XT index v2 (Illumina #FC-131-1002). The index PCR products were cleaned with the Agencourt AMPure XP kit (Beckman Coulter #A63881) and checked for quality with a High Sensitivity TapeStation chip. Cleaned libraries were quantified using a Qubit 4.0 system with High-Sensitivity Kit (ThermoFisher Scientific), and 28 nM of each sample was pooled.

### Zaprinast, LPA and *E. coli* OMVs treatments

Explant form *Gpr35^WT^* mice were pretreated with either 10 μM Zaprinast or LPA for 1.5h followed by 10 μg of *E. coli* OMVs for 1.5h.

### Quantification and Statistical Analysis

Data are presented as dot plots of individual values with medians. Statistical significance analysis was performed with GraphPad Prism software; either Mann-Whitney U, or two-way ANOVA tests were performed depending on the experimental setting. Data was further analyzed with Grubbs’ test to identify the outliers. Differences were considered significant as follows: ^∗^p < 0.05, ^∗∗^p < 0.01, ^∗∗∗^p < 0.001.

## Acknowledgments

We thank Silvia Kobel and Aria Minder from the Genomics Facility Basel, Department of Environmental Systems Science, ETH Zurich, for helping with the scRNA-seq. We also thank the Histology Facility Basel, University of Basel, for helping with the IHC staining and the 16S *in situ* hybridization. Calculations were performed at sciCORE (http://scicore.unibas.ch/) scientific computing center at the University of Basel. The SNSF grant 310030_175548 to J.H.N. financed the publication of our manuscript without restrictions. The SNSF M.D. Ph.D. fellowship 323530_183981 supported T.K.

## Author Contributions

H.M, B.K, T.K, P.W, M.L.B, R.A.M performed experiments. J.R performed bioinformatics analysis of the scRNA-seq data. C.C.W provided the VillinCre mice. J.C.W, performed bioinformatics analysis of the 16S rRNA-seq data. C.U.R, provided the *C. rodentium*. B.K, T.K, P.W, M.L.B, J.C.W, R.A.M, E.F, J.R, P.L, P.H, and E.J.V, corrected the paper. H.M and J.H.N designed the study and wrote the paper. J.H.N. conceived the idea. All authors discussed the data and read and approved the manuscript.

## Supplementary figures and tables

**Figure S1.**
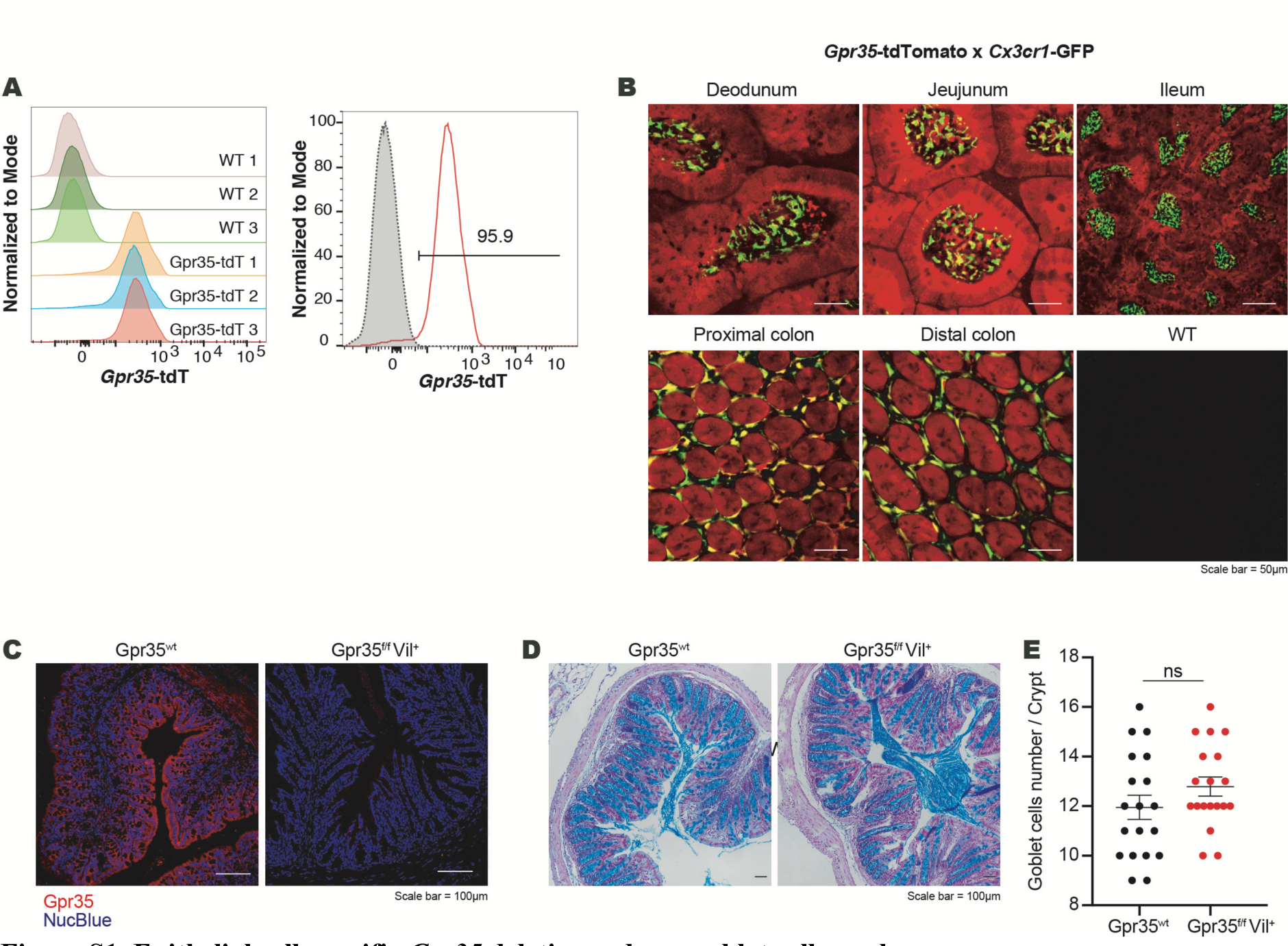
Epithelial cell-specific *Gpr35* deletion reduces goblet cell numbers. (A) Representative *Gpr35*-tdTomato expression by flow cytometry in epithelial cells (Epcam^+^, CD45^-^) from proximal colon of *Gpr35*-tdTomato reporter mice (red unfilled histograms) and *Gpr35^wt^* mice (gray histograms) as the background control. (B) *Ex vivo* fluorescence imaging of duodenum, jejunum, ileum, proximal colon and distal colon, from *Gpr35*- tdTomato (red) x *Cx3cr1*-GFP (green) double reporter mice. The last panel shows colon from WT as the background control. (C) Confocal immunofluorescence images of proximal colon sections obtained from *Gpr35^f/f^Vil^+^* and *Gpr35^wt^* littermates and stained for Gpr35 (red) and DAPI (blue). Scale bars, 100 µm. (D) Representative AB/PAS staining of distal colon sections obtained from *Gpr35^f/f^Vil^+^* and *Gpr35^wt^* littermates. Scale bars, 50 µm. (E) Cell count of goblet cells in (D) performed blindly by two different investigators in at least 30 crypts. Data are represented as mean ± SEM ns, not significant by Mann-Whitney test.

**Figure S2.**
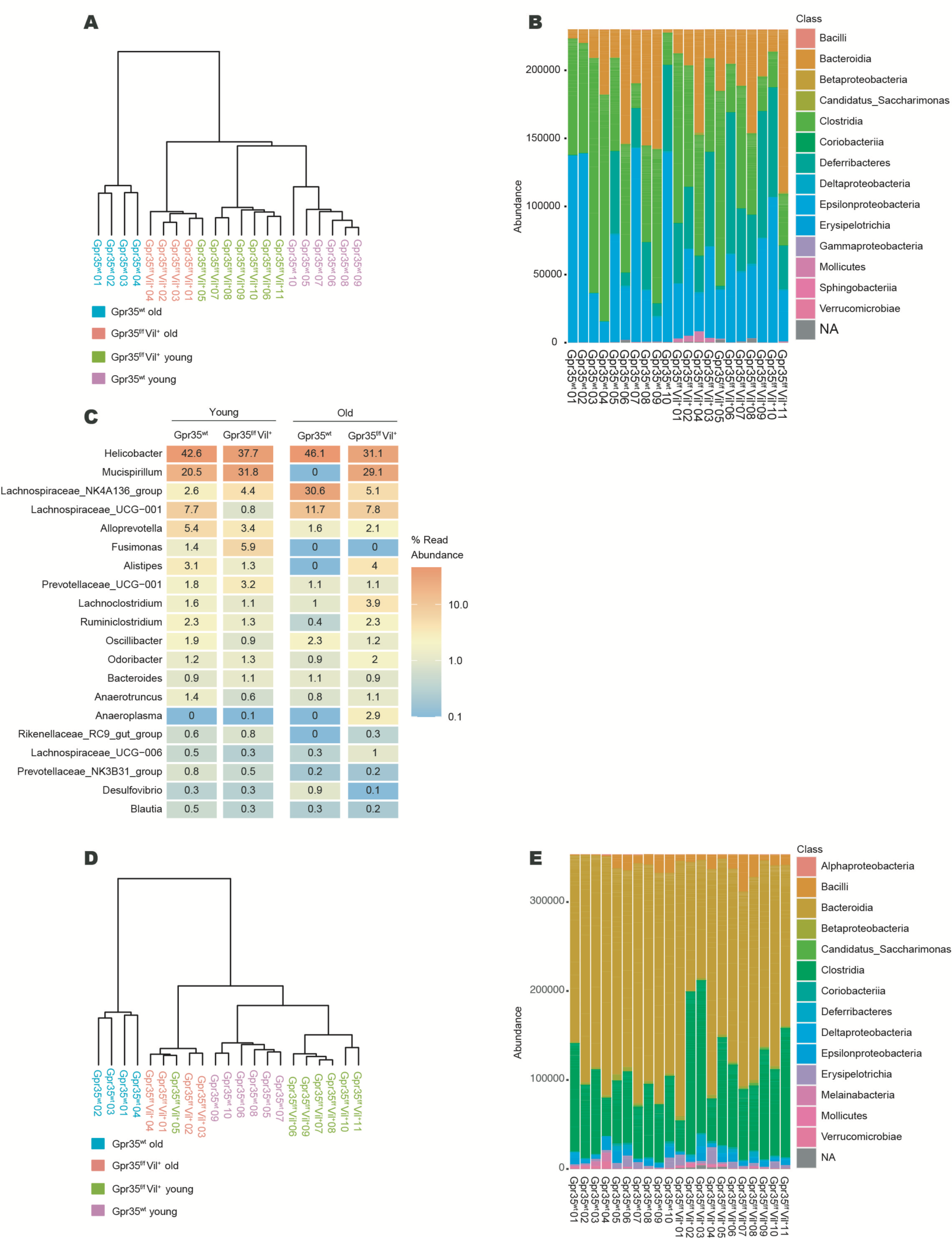
Epithelial cell-specific *Gpr35* deletion correlates with an altered mucosa-associated microbiome. (A) Hierarchical clustering for mucosal bacterial samples based on Bray-Curtis dissimilarity and Ward’s minimum variance method (ward.D2). (B) Relative abundance of mucosal-bacterial communities at the class level in *Gpr35^f/f^Vil^+^* and *Gpr35^wt^* at different age points. (C) Relative abundance of taxonomic groups at the genus level averaged across mucosa-associated bacteria samples of old and young *Gpr35^f/f^Vil^+^* and *Gpr35^wt^* littermates. (D) Hierarchical clustering for fecal bacterial samples based on Bray-Curtis dissimilarity and Ward’s minimum variance method (ward.D2). (E) Relative abundance of fecal-bacterial communities at the class level in *Gpr35^f/f^Vil^+^* and *Gpr35^wt^* at different age points.

**Figure S3.**
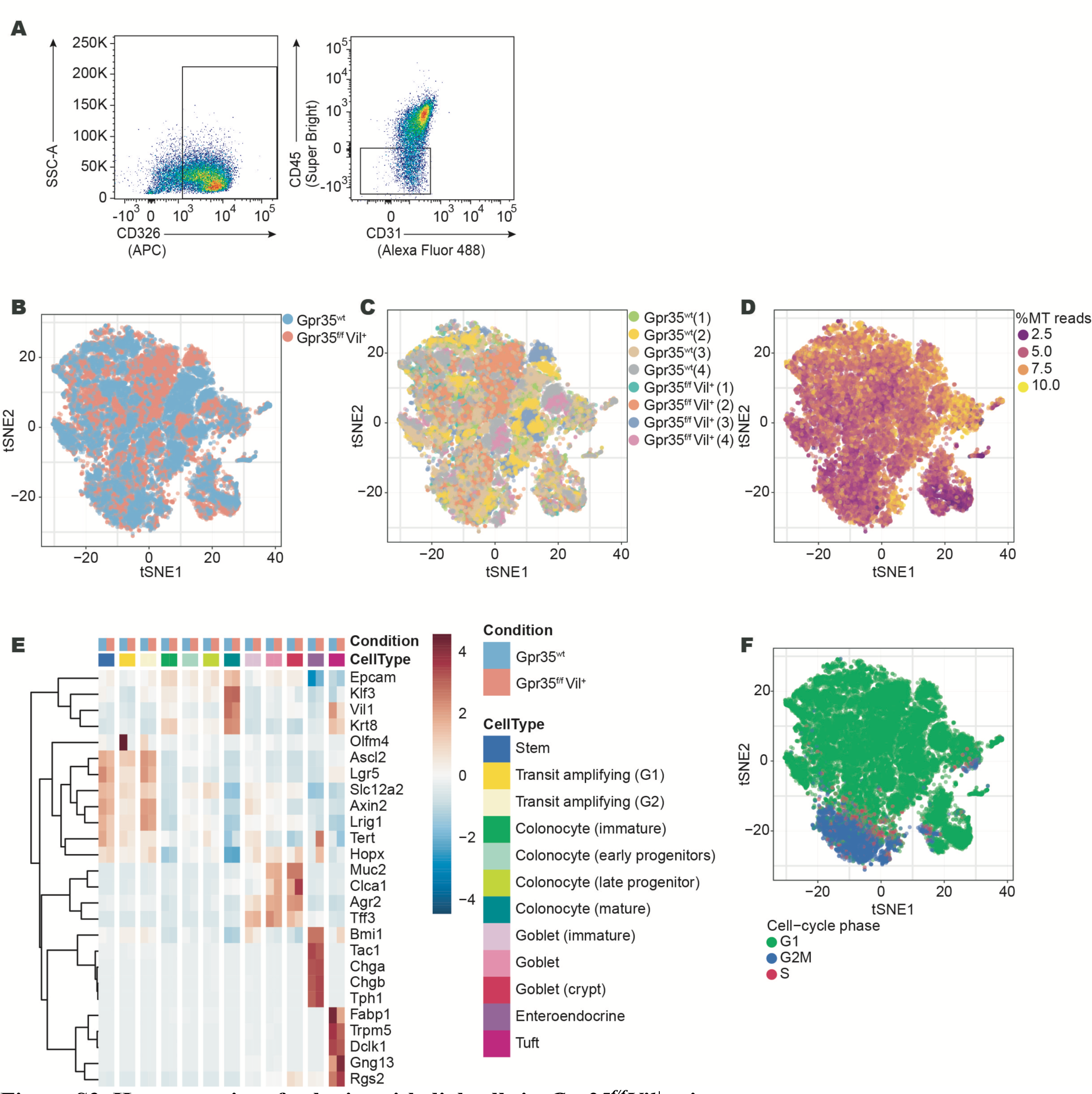
Heterogeneity of colonic epithelial cells in *Gpr35^f/f^Vil^+^* mice. (A) Flow-cytometry analysis of cells isolated form proximal colon of *Gpr35^f/f^Vil^+^* and *Gpr35^wt^* littermates before scRNA-seq. (B-D, F) *t*-SNE plot showing proximal colonic epithelial cells from *Gpr35^f/f^Vil^+^*(n = 4) and *Gpr35^wt^* (n = 4) littermates assayed via scRNA-seq, and highlighting: (B) the distribution of *Gpr35^f/f^Vil^+^* and *Gpr35^wt^* cells; (C) the distribution of cells from individual mice replicates; (D) the percentage of reads originating from mitochondrial genes; (F) the inferred cell-cycle phase. (E) Heatmap showing the centered and scaled average expression across cell types and conditions of known cell type marker genes.

**Figure S4.**
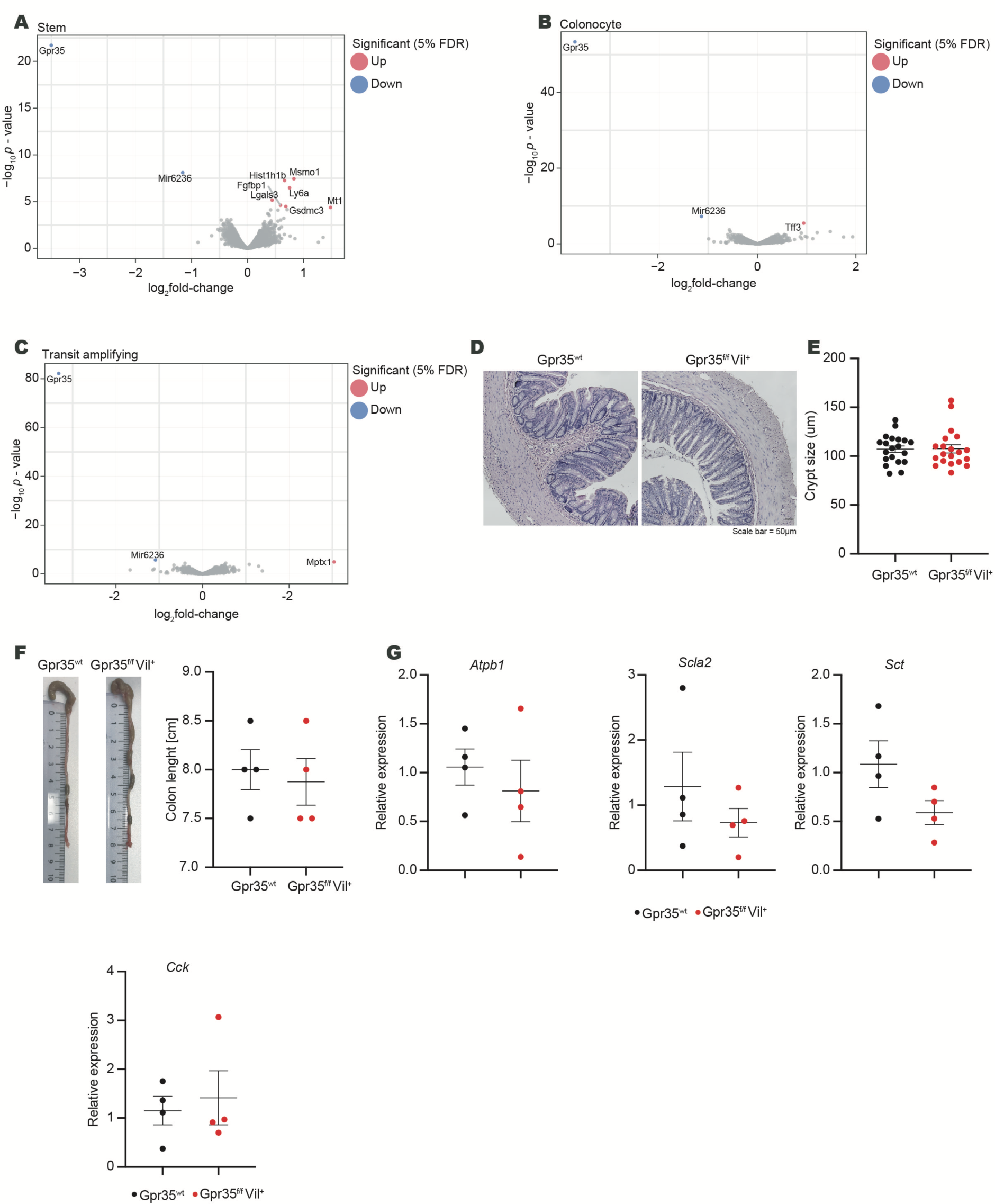
Increased pyroptotic signatures in goblet cells lacking epithelial *Gpr35*. (A-C) Volcano plot showing genes differentially expressed between *Gpr35^f/f^Vil^+^* (n = 4) and *Gpr35^wt^* (n = 4) (A) stem cells, (B) colonocytes or (C) transit amplifying cells. Legend similar to Figure 4A. (D) Representative H&E images of proximal colon sections obtained from *Gpr35^f/f^Vil^+^* and *Gpr35^wt^* littermates. (E) Crypt size quantified from (D). (F) Colon lengths of 10 week old *Gpr35^f/f^Vil^+^* and *Gpr35^wt^* mice. (G) mRNA expression levels of *Atpb1, Scla2, Sct and Cck* measured by qRT-PCR in proximal colon samples obtained from *Gpr35^f/f^Vil^+^* (n = 4) and *Gpr35^wt^* littermates (n = 4). Each dot represents one animal with medians. Data are represented as mean ± SEM *p ≤ 0.05, **p ≤ 0.01, ***p ≤ 0.001, ****p ≤ 0.0001 by Mann-Whitney in (E), (F) and (G).

**Figure S5.**
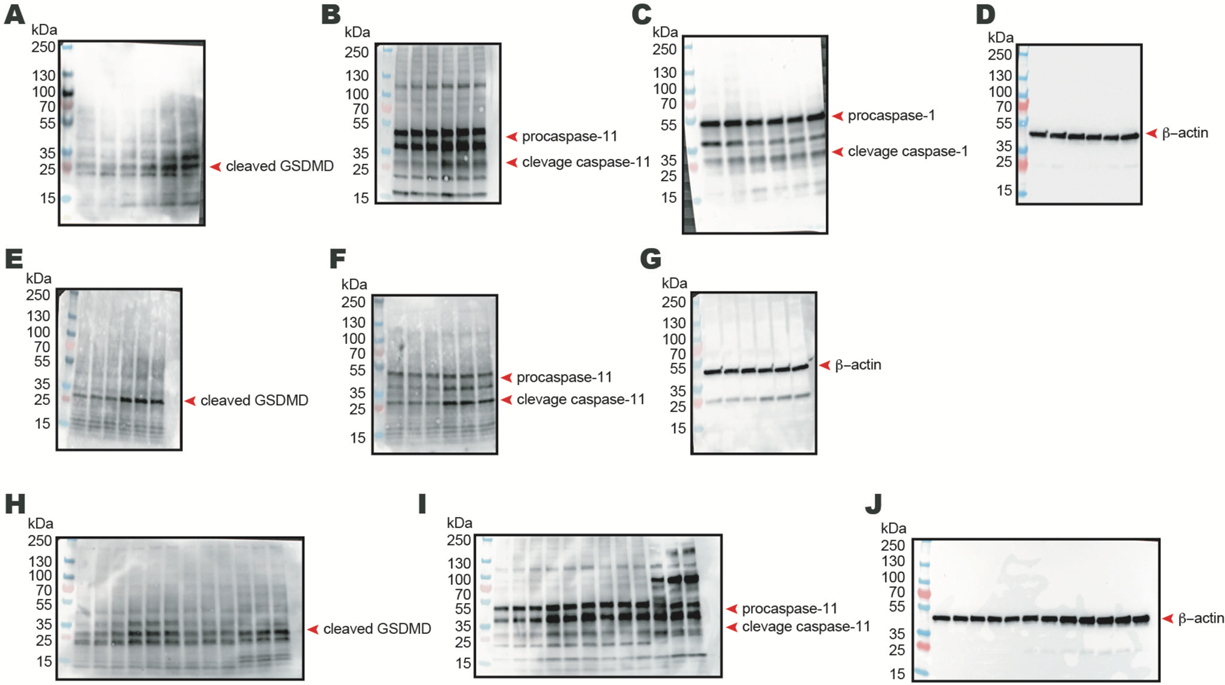
Pyroptosis upon *Gpr35* deletion is caspase-11 dependent. (A-J) Whole membrane images of western blots represented in Figure 5A-5F.

**Figure S6.**
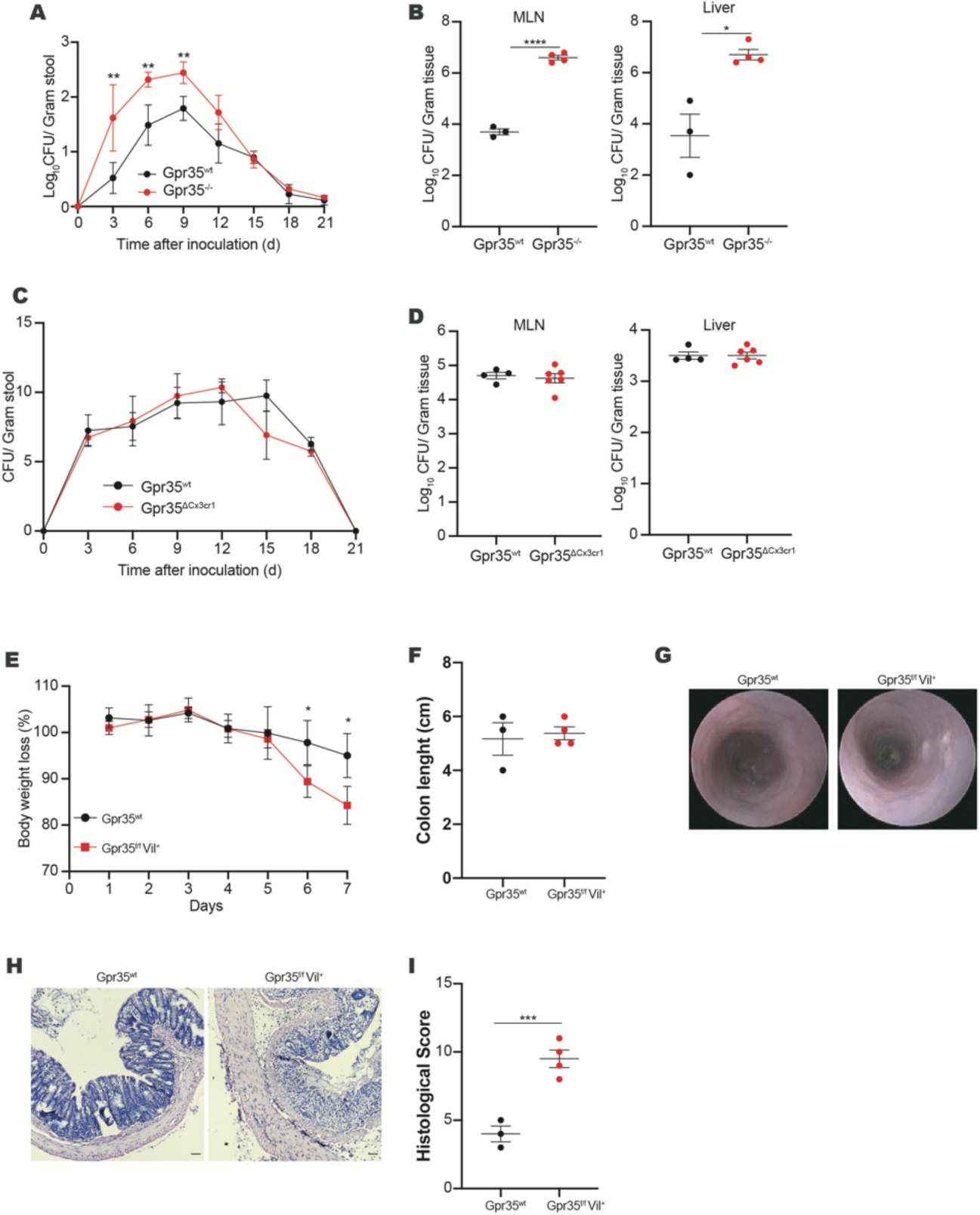
Epithelial GPR35 protects against *Citrobacter rodentium* infection. (A and B) *C. rodentium* CFU/g of (A) feces, (B) MLN and liver from *Gpr35^-/-^* (n = 4) and *Gpr35^wt^* (n = 3) littermates. (C and D) *C. rodentium* CFU/g of (C) feces, (D) MLN and liver from *Gpr35^ΔCx3cr1^* (n = 6) and *Gpr35^wt^* (n = 4) littermates. (E) Body weight changes of *Gpr35^f/f^Vil^+^* (n = 4) and *Gpr35^wt^* (n = 3) littermates (normalized to initial weight). (F) Colon lengths of *Gpr35^f/f^Vil^+^* and *Gpr35^wt^* mice on day 7. (G) Endoscopic images of *Gpr35^f/f^Vil^+^* and *Gpr35^wt^* mice. (H) Representative H&E images of proximal colon sections obtained from *Gpr35^f/f^Vil^+^* and *Gpr35^wt^* littermates. (I) Histology scores quantified from (H). Each dot represents one animal with medians. Data are represented as mean ± SEM *p ≤ 0.05, **p ≤ 0.01, ***p ≤ 0.001, ****p ≤ 0.0001 by two-way ANOVA with Tukey’s multiple comparisons test in (A), (C) and (E) or Mann-Whitney in (B), (D), (F) and (I).

Table S1: Genes differentially expressed in each cell type of Gpr35^f/f^Vil^+^ and Gpr35^wt^ littermates. (See separated uploaded excel file).

**Table S2:**
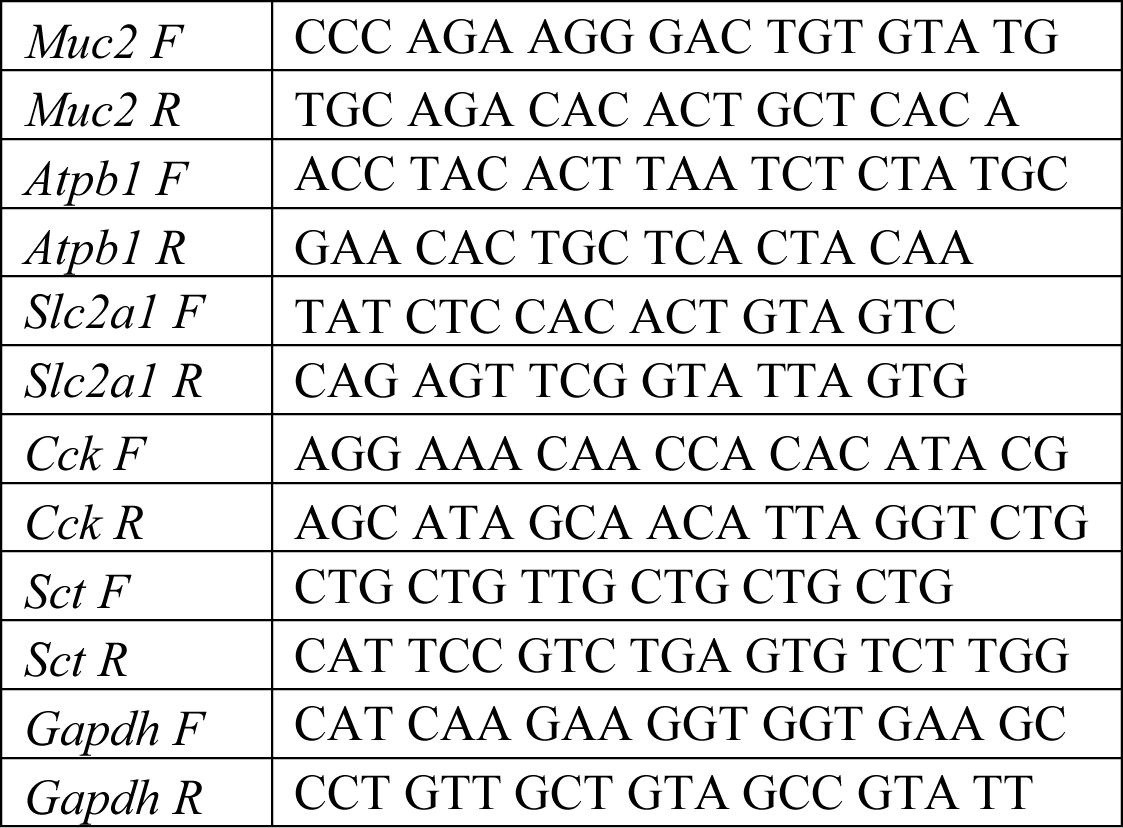
q-PCR primers.

**Table S3:** 16S primers.

|  |  |
| --- | --- |
| Forward | GTG**NC**AGCMGCCGCGGTAA |
| Reverse | GGACTACHVGGGT**NT**CTAA |

